# Multipotent progenitors and hematopoietic stem cells arise independently during the endothelial to hematopoietic transition in the early mouse embryo

**DOI:** 10.1101/2021.04.13.439736

**Authors:** Tessa Dignum, Barbara Varnum-Finney, Sanjay Srivatsan, Stacey Dozono, Olivia Waltner, Adam Heck, Cynthia Nourigat-McKay, Dana L. Jackson, Shahin Rafii, Cole Trapnell, Irwin D. Bernstein, Brandon Hadland

## Abstract

During embryogenesis, waves of hematopoietic progenitors develop from hemogenic endothelium (HE) prior to the emergence of self-renewing hematopoietic stem cells (HSC). Although previous studies have shown that yolk sac-derived erythromyeloid progenitors and HSC emerge from distinct populations of HE, it remains unknown whether the earliest lymphoid-competent progenitors, multipotent progenitors, and HSC originate from common HE. Here we demonstrate by clonal assays and single cell transcriptomics that rare HE with functional HSC potential in the early murine embryo are distinct from more abundant HE with multilineage hematopoietic potential that fail to generate HSC. Specifically, HSC-competent HE are characterized by expression of CXCR4 surface marker and by higher expression of genes tied to arterial programs regulating HSC dormancy and self-renewal. Together, these findings suggest a revised model of developmental hematopoiesis in which the initial populations of multipotent progenitors and HSC arise independently from HE with distinct phenotypic and transcriptional properties.

## INTRODUCTION

Hematopoietic stem cells (HSC) possess simultaneous multilineage and self-renewal capacity and are defined by their ability to reconstitute the entire hematopoietic system upon transplantation to a conditioned host. HSC emerge during embryonic development from a specialized subset of endothelial cells (EC) with hematopoietic potential, known as hemogenic endothelium (HE). This endothelial-to-hematopoietic transition (EHT) produces the first transplantable HSC by murine embryonic day 10.5-11 (E10.5-11) within arterial vessels including the aorta of the para-aortic-splanchnopleura/aorta-gonad-mesonephros region (P-Sp/AGM) and the vitelline and umbilical arteries (de Bruijn et al., 2000; Medvinsky and Dzierzak, 1996; Müller et al., 1994). Prior to the emergence of HSC, however, several distinct waves of hematopoiesis provide the developing embryo with its initial functioning blood cells. The first of these, referred to as the primitive and erythromyeloid progenitor (EMP) waves, emerge in the yolk sac and are primarily restricted to the production of erythroid, megakaryocytic, and myeloid progeny (Palis, 2016). These waves lack significant lymphoid potential, which emerges during a third wave of hematopoiesis that produces lympho-myeloid progenitors (LMP), multipotent progenitors (MPP), and ultimately HSC (Böiers et al., 2013; Hadland and Yoshimoto, 2018; Inlay et al., 2014). Since clonal MPP are detectable prior to HSC emergence, it has been postulated that they give rise to HSC following their acquisition of additional properties necessary for HSC function, such as self-renewal (Inlay *et al*., 2014). However, this remains to be formally determined, as initial MPP activity has not been clonally linked to later HSC fate, and surface markers that can prospectively distinguish HE with HSC potential from those generating various categories of multilineage progenitors have not been identified. To explore this, we employed single cell index sorting of hemogenic precursors to co-culture with AGM-derived endothelial cells (AGM-ECs) and assays for HSC and multilineage potential. We previously reported that this co-culture system supports the formation of multilineage-engrafting HSC from P-Sp/AGM-derived HE isolated as early as E9 (Hadland et al., 2017; Hadland et al., 2018; Hadland et al., 2015). Focusing our analysis on HE clonally isolated from E9-E10 P-Sp/AGM at the onset of EHT, we found that CXCR4 expression distinguishes HSC-competent HE from progenitor-restricted HE, including a subset with in vitro multilineage potential. Single-cell RNA sequencing (scRNAseq) demonstrated that *Cxcr4*-expressing HE are simultaneously enriched in expression of arterial-specific genes and HSC self-renewal genes controlling dormancy and cell cycle, compared to the more abundant *Cxcr4*-negative HE. Altogether, our studies suggest a model in which the earliest MPP and HSC emerge independently from phenotypically and transcriptionally distinct populations of HE, which has important implications for longstanding efforts to generate HSC from pluripotent stem cells.

## RESULTS

### CXCR4 expression marks rare clonal HE with HSC potential in the early embryo

Using AGM-EC co-culture and transplantation assays, we previously demonstrated that long-term HSC activity within the P-Sp/AGM-derived VE-Cadherin^+^ population between E9 and E11.5 is largely restricted to cells co-expressing EPCR and CD61 (henceforth referred to as the V^+^E^+^61^+^ population) (Hadland et al., 2021; Hadland *et al*., 2017; Hadland *et al*., 2018). Employing our previously described method of single-cell index sorting to AGM-EC co-culture followed by immunophenotypic analysis of HSC activity—defined by VE-Cadherin^-/lo^Gr1^-^F4/80^-^Sca-1^+^EPCR^hi^ phenotype, a reliable surrogate for multilineage engraftment potential in this assay (Hadland *et al*., 2017; Hadland *et al*., 2018)—we determined that the majority of clonal V^+^E^+^61^+^ hemogenic precursors (defined here as CFC, colony-forming cells) isolated between E9 and E10 generate hematopoietic progenitor cells (HPC-CFC) lacking HSC activity, whereas those with HSC potential (HSC-CFC) are rare (Figure 1A-B). If HE with cell-intrinsic HSC potential constitute a distinct population from HE that lack HSC potential, we hypothesized that they may be prospectively isolated based on expression of unique surface markers. Previous studies have shown that arterial markers DLL4 and CD44 are expressed by HE in the P-Sp/AGM, but do not necessarily distinguish HE with HSC potential from those restricted to progenitor fates at this early stage (Hadland *et al*., 2017; Oatley et al., 2020). We hypothesized that another arterial marker, CXCR4, may better predict HSC potential given our recent observation that its canonical ligand, CXCL12, is required for the generation of HSC (but not progenitors) from P-Sp/AGM-derived hemogenic precursors in a stroma-free engineered niche (Hadland *et al*., 2021), and that CXCR4 is heterogeneously expressed within the V^+^E^+^61^+^ population of the E9.5 P-Sp/AGM which is enriched in arterial markers DLL4 and CD44 (Figure 1C).

**Figure 1.**
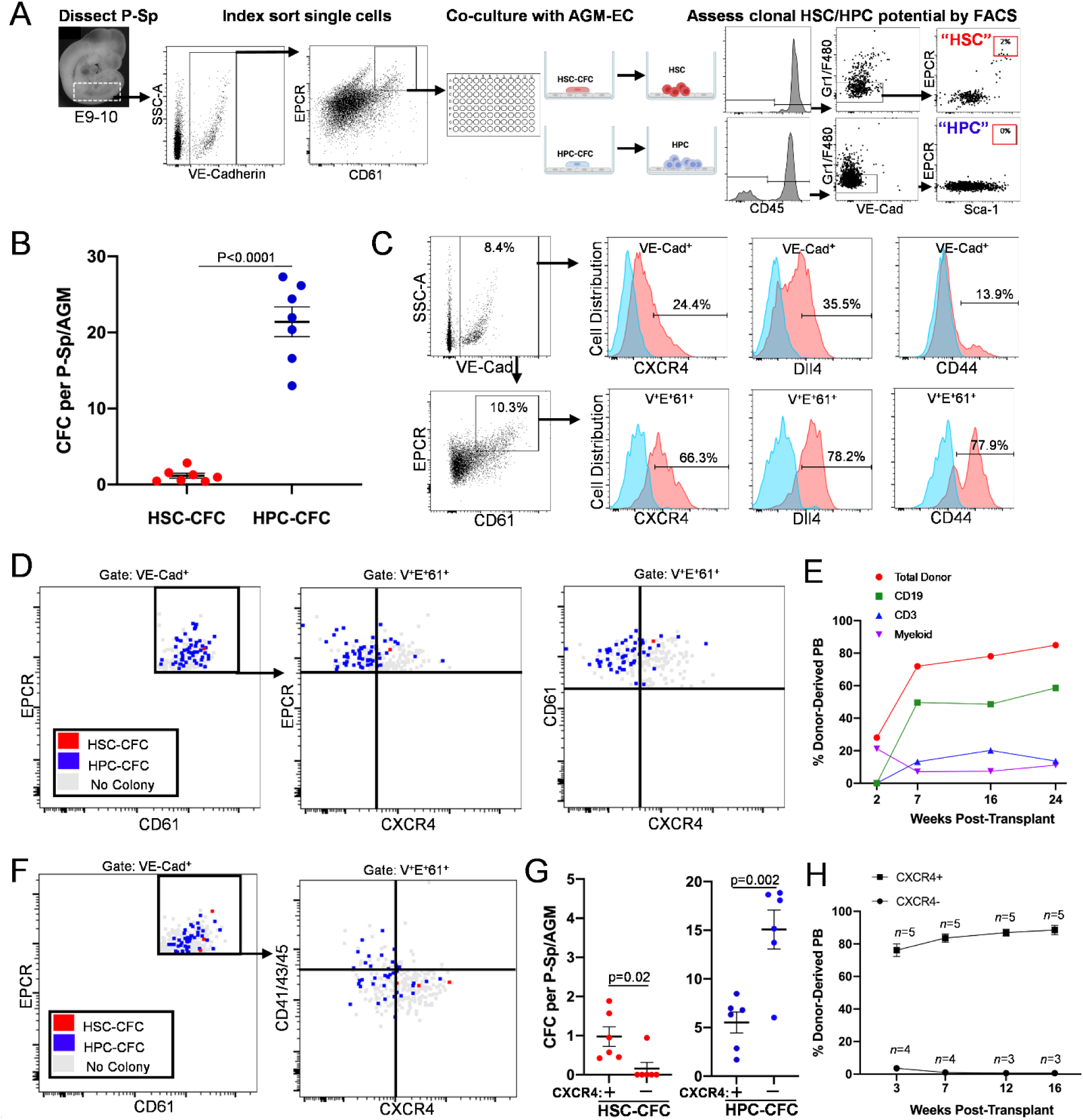
CXCR4 expression marks rare clonal HE with HSC potential in the early murine embryo. **A.** Methodology for index sorting of single P-Sp-derived V^+^E^+^61^+^ cells to co-culture on AGM-EC and subsequent analysis to assess HSC potential detected by immunophenotype (CD45^+^VE-Cadherin^-/lo^Gr1^-^F4/80^-^Sca-1^+^EPCR^hi^). **B.** HSC colony-forming cells (HSC-CFC) and HPC colony-forming cells (HPC-CFC) detected across 7 independent experiments from E9-E10 (19-30 sp). Error bars show mean +/− SEM. **C.** Expression of CXCR4, CD44, and DLL4 in the VE-Cadherin^+^ population and the V^+^E^+^61^+^ subset of E9.5 (29 sp) P-Sp. Blue histograms indicate isotype controls. **D.** Correlation of clonal HSC-CFC and HPC-CFC potential with expression CXCR4 on individual index-sorted V^+^E^+^61^+^ cells at E9.5 (26-29 sp). **E.** Donor-derived peripheral blood (PB) engraftment of clonal progeny of HSC-CFC in (D), including B lymphoid (CD19), T lymphoid (CD3), and myeloid contribution. **F.** Correlation of clonal HSC-CFC and HPC-CFC with surface expression of CXCR4 and CD41, CD43, and CD45 on individual index-sorted V^+^E^+^61^+^ cells at E9.5 (29-30 sp). **G**. Distribution of CXCR4^+^ and CXCR4^-^ HSC-CFC and HPC-CFC detected across 6 independent experiments from E9-E10 (19-30 sp). Error bars show mean +/− SEM. **H.** Donor-derived PB engraftment of bulk-sorted CXCR4^+^ and CXCR4^-^ cells within the V^+^E^+^ population at E9.5 (22-29 sp). Error bars show mean +/− SEM; *n* indicates number of recipient mice. All recipients of CXCR4^+^ cells had multilineage engraftment (See also Figure S1, Table S1).

To investigate whether expression of CXCR4 distinguishes clonal HSC precursors, we isolated V^+^E^+^61^+^ cells from E9-E10 P-Sp/AGM for single-cell index sorting to AGM-EC co-culture (Figure 1A). Correlating phenotypic HSC output following co-culture with the initial index phenotype of each V^+^E^+^61^+^ starting cell, we show that the rare hemogenic precursors (including CD41^-^CD43^-^CD45^-^ HE) with HSC potential (HSC-CFC) typically express CXCR4, whereas the majority of HPC-CFC lack CXCR4 expression (Figure 1D, F, G, Figure S1A, B). Based on our previous studies of clonal HSC precursors at this early stage, HSC-CFC as defined by immunophenotype can be heterogenous in regards to long-term multilineage engraftment in vivo (Hadland *et al*., 2017). Thus, to validate clonal CXCR4^+^ HSC-CSF included functional long-term HSC, we assessed a subset of the HSC-by-phenotype colonies through transplant and confirmed their multilineage engraftment potential in vivo (Figure 1E, Supplementary Table 1A). In addition, when we sorted and separately transplanted the CXCR4^+^ and CXCR4^-^ fractions of the E9-9.5 P-Sp/AGM-derived V^+^E^+^ population in bulk (Figure S1C, D) we confirmed that HSC potential was confined to the CXCR4^+^ subset, whereas the CXCR4^-^ fraction generated phenotypic HPC that lacked significant multilineage engraftment (Figure 1H, Figure S1E, Supplementary Table 1B). Altogether, these findings show that CXCR4 expression marks rare HSC-competent HE in the early embryo.

### Clonal HE harboring multilineage hematopoietic activity emerge independently from HSC-competent HE in the early embryo

Although the majority of E9-E10 P-Sp/AGM-derived V^+^E^+^61^+^ HE generate HPC lacking multilineage engraftment potential, the lineage potentials of clonal HPC-CFC within this population remain undefined. In accordance with recent literature, we hypothesized that HPC-CFC detected in our co-culture system are precursors to the initial wave of LMP that are proposed to be HSC-independent (Böiers *et al*., 2013; Zhu et al., 2020). Clonal precursors with in vitro multilineage hematopoietic potential have also been identified in the E9-10 yolk sac and P-Sp/AGM (Inlay *et al*., 2014), and were postulated to represent HSC precursors that acquire definitive HSC fate upon further maturation (thus equivalent to HSC-CFC in our assays).

To investigate this, we explored the in vitro lineage potentials of clonal E9-10 HPC-CFC following AGM-EC co-culture by secondary colony-forming unit (CFU) assays (to screen for erythroid and myeloid potential) and co-culture with OP9 or OP9-DLL4 stromal cells (to assess B- and T-lymphoid potential, respectively) (Figure 2A). Notably, we observed a variety of lineage-restricted potentials beyond LMP, including erythromyeloid and myeloid-only, from HPC-CFC expressing hematopoietic markers CD41, CD43, and/or CD45, as well as from HPC-CFC with HE phenotype (CD41^-^CD43^-^ CD45^-^) and HPC-CFC expressing arterial marker CD44 (Oatley *et al*., 2020) (Figure 2B-D, Figure S2A, B). We also detected HPC-CFC with multipotent hematopoietic potential (MPP-CFC) that are mostly CXCR4^-^, are more abundant than HSC-CFC, and fail to generate phenotypic or functionally engrafting HSC following AGM-EC culture (Figure 2B-D, Figure S2A, B, and data not shown). These data demonstrate the unexpected existence of a unique population of MPP-competent HE that lack cell-intrinsic properties necessary for HSC fate.

**Figure 2.**
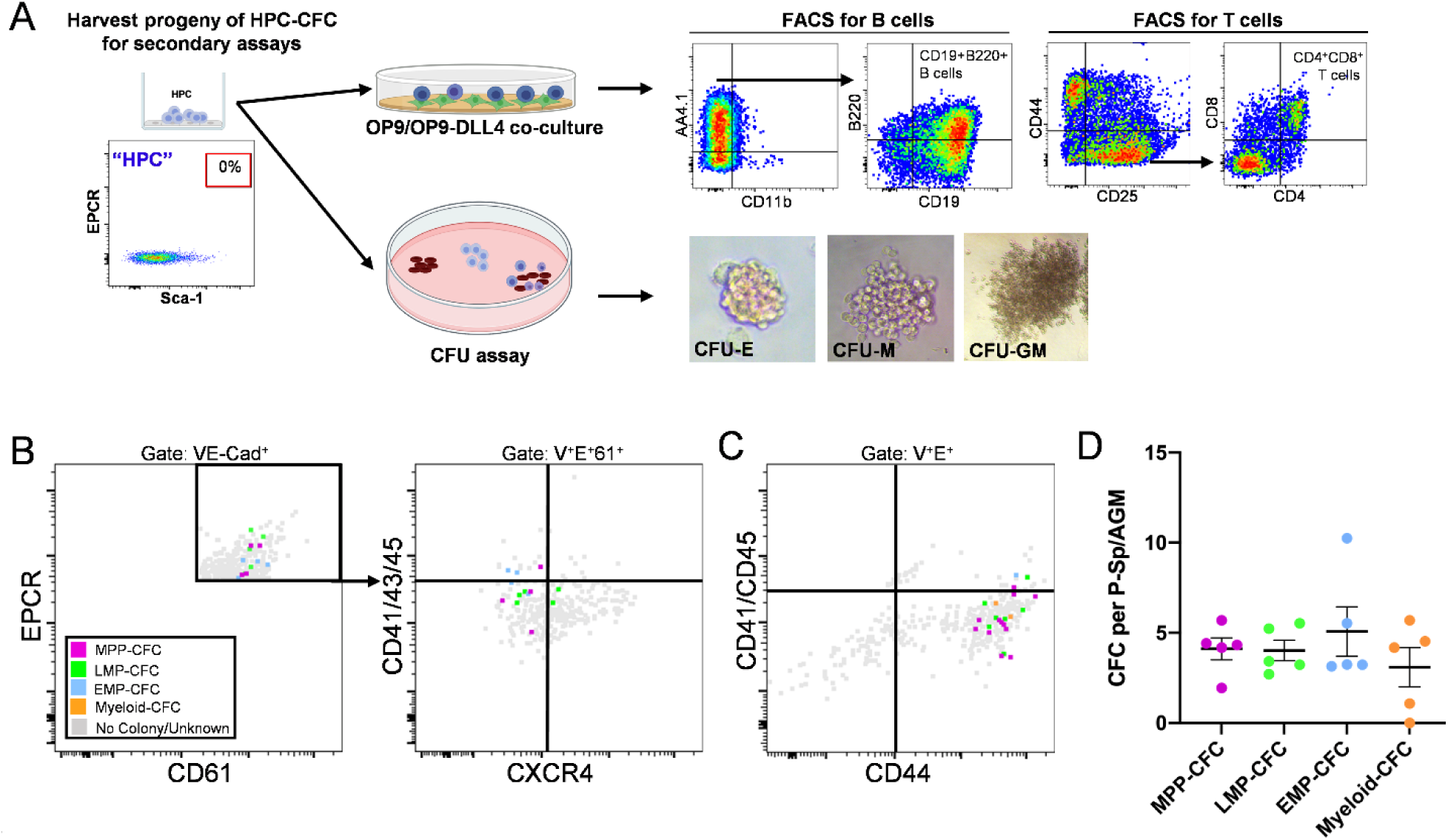
Clonal HE harboring multilineage hematopoietic activity emerge independently from HSC-competent HE in the early embryo. **A.** Methodology for analysis of in vitro hematopoietic lineage potential of clonal progeny of HPC-CFC. Representative images of erythroid colony forming unit (CFU-E) (20X), macrophage colony forming unit (CFU-M) (20X), and granulocyte-monocyte colony forming unit (CFU-GM) (4X) shown. **B-C.** Correlation of clonal hematopoietic lineage potential with CXCR4, CD41, CD43, CD44, and CD45 expression of individual HPC-CFC from E9.5 P-Sp (26-28 sp). Multipotent (MPP-CFC), lymphomyeloid (LMP-CFC), erythromyeloid (EMP-CFC), and myeloid colony-forming cells (Myeloid-CFC). No colony/unknown (gray) indicates absence of detectable hematopoietic output following AGM-EC or secondary CFU/OP9 assays. **D.** Total progenitor CFC within the V^+^E^+^61^+^ population per P-Sp detected across 5 independent experiments from E9 to E10 (19-30 sp). Error bars show mean +/− SEM. (See also Figure S2).

### Single cell RNA-sequencing identifies the transcriptional signature of *Cxcr4*-expressing HSC-competent HE

Based on our finding that CXCR4 expression identifies rare clonal HSC-competent HE in the V^+^E^+^61^+^ population, we leveraged single cell transcriptomics to investigate the transcriptional landscape of hemogenic precursors differentially expressing *Cxcr4*, hypothesizing that *Cxcr4* expression would reveal the unique transcriptional state of HE with HSC potential. To this end, we dissected P-Sp from E9-9.5 embryos and sorted V^+^E^+^61^+^ cells for scRNAseq (Figure S3A-B). Using the Monocle3 analysis toolkit (Cao et al., 2019), we performed dimensionality reduction by UMAP and unsupervised clustering of the single cell transcriptomes (Figure 3A, Figure S3C-D). After removing clusters expressing somite-specific genes (presumed to represent a population of contaminating cells detected in the post-sort analysis, Figure S3A, E), cells in the remaining clusters expressed EC marker *Cdh5* (VE-Cadherin) and were heterogeneous for genes associated with arterial EC and HE (Figure 3B), consistent with immunophenotypic and functional analysis of V^+^E^+^61^+^ cells (Figure 1-2). A small cluster (23) expressing markers of mature HPC (*Ptprc/*CD45, *Flt3*, etc) was also identified (Figure 3B, Figure S3F). To identify clusters associated with *Cxcr4* expression, we classified the clusters in UMAP space based on threshold detection of *Cxcr4* transcript in single cells in each cluster (Figure 3C). We then classified cells according to type on the basis of known marker genes using Garnett (Pliner et al., 2019), assigning cells as unspecified EC, early and mature arterial EC, HE, and HPC (Figure 3D). Cxcr4-negative clusters were enriched in unspecified EC, early arterial EC, and HPC, whereas Cxcr4-positive clusters were enriched in mature arterial EC (Figure 3D). Both clusters contained HE, with fewer in the Cxcr4-positive clusters relative to the Cxcr4-negative clusters (Figure 3D), which is consistent with the relative frequencies of clonal CFC between CXCR4^+^ and CXCR4^-^ subsets of the V^+^61^+^E^+^ population in our single cell index assays (Figure 1D, F, Figure S1A, B). We next examined the relative aggregate expression (gene-set score) of published arterial EC-specific genes (Aranguren et al., 2013; Luo et al., 2021; Xu et al., 2018). As expected, these gene-set scores were highest in cells transcriptionally classified as mature arterial EC. Furthermore, Cxcr4-positive HE had significantly higher arterial EC gene-set scores than Cxcr4-negative HE (Figure 3E, Figure S3G). We also examined previously published HSC-specific signature gene sets from E11 AGM (Vink et al., 2020) and adult bone marrow (Cabezas-Wallscheid et al., 2017; Chambers et al., 2007; Rodriguez-Fraticelli et al., 2020; Wilson et al., 2015). Remarkably, we found that these gene-set scores followed a similar pattern to that of arterial EC-specific genes (peaking in mature arterial EC), and were significantly higher in Cxcr4-positive HE than in Cxcr4-negative HE (Figure 3E, Figure S3G). When we examined expression of genes recently reported as specific to HSC-primed HE in the P-Sp/AGM (Hou et al., 2020), this gene-set score peaked in HE cell type in our data, as expected. However, it was significantly higher in Cxcr4-negative compared with Cxcr4-positive HE, suggesting that this gene set is not necessarily specific to HE with HSC competence at this early stage, but is also expressed by HE restricted to multilineage progenitor fates (Figure 3E).

**Figure 3.**
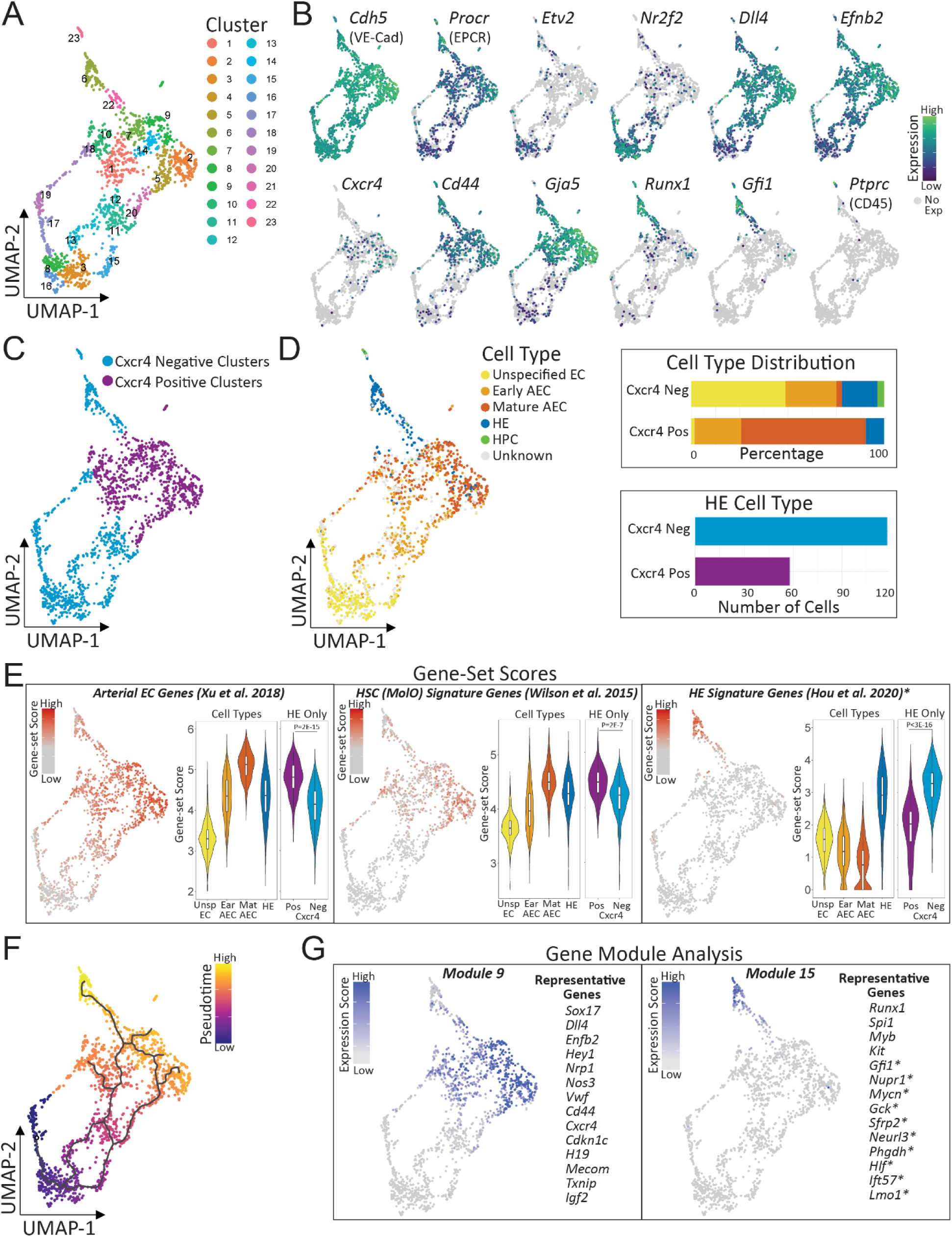
Single cell RNA-sequencing identifies the transcriptional signature of *Cxcr4*-expressing HSC-competent HE. **A.** UMAP and cluster analysis of scRNAseq data from E9-9.5 P-Sp-derived V^+^E^+^61^+^ cells. **B.** Gene expression heatmap for pan-endothelial marker VE-Cadherin (*Cdh5*), EPCR (*Procr*), *Cxcr4*, and selected transcripts used for cell type classification in (D). **C**. Classification of clusters based on *Cxcr4* transcript detected in >5% of cells in each cluster from (A). **D.** Left: Cell type classification by Garnett based on marker genes (see methods for marker genes). Right: Distribution of cell types among Cxcr4+ and Cxcr4-clusters and quantification of HE cell type within Cxcr4+ and Cxcr4-clusters. **E.** Left: Heatmap of gene-set scores for signature gene sets defining arterial EC (Xu *et al*., 2018), HSC (Wilson *et al*., 2015), and HE (Hou *et al*., 2020). Right: Violin plots of gene-set scores by cell type, and by Cxcr4+ vs Cxcr4-HE. P values indicate Wilcoxon rank-sum test. **F.** Pseudotime trajectory. **G.** Heatmaps for expression scores of gene modules that co-vary over pseudotime. Gene modules 9 and 15 are depicted with selected subset of representative genes from each (See also Figure S3, Table S3).

As a complementary approach to identify genes enriched in the Cxcr4-positive clusters in our scRNAseq data, we next determined modules of genes that vary as a function of pseudotime (Cao *et al*., 2019; Trapnell et al., 2014) (Figure 3F). This analysis revealed two unique gene modules (9 and 15) whose expression patterns correlate with regions of UMAP containing Cxcr4-positive and Cxcr4-negative HE, respectively (Figure 3G, Figure S3H). Significantly upregulated genes in module 9 include *Cxcr4,* along with arterial EC-specific genes and several genes required for HSC self-renewal (including *H19*, *Mecom*, *Cdkn1c*, *Igf2*, and *Txnip*) (Jeong et al., 2009; Kataoka et al., 2011; Matsumoto et al., 2011; Thomas et al., 2016; Venkatraman et al., 2013) (Figure 3G, left panel, and Table S2). In contrast, module 15 is enriched in genes associated with EHT and definitive hematopoiesis—including, notably, 10 of the 11 HE-specific signature genes identified by Hou *et al*. (2020) (Figure 3G, right panel and Table S2).

Finally, we applied regression analysis to identify genes that are differentially expressed between Cxcr4-positive and Cxcr4-negative HE (adjusted p value <0.05, Table S4). Gene ontology analysis identified enrichment of genes in Cxcr4-positive HE associated with angiogenesis, cell motility, and adhesion, whereas Cxr4-negative HE were enriched in genes associated with protein synthesis, metabolic activity, and proliferation (Figure S4A, B, Table S4). Remarkably, genes enriched in Cxcr4-positive HE showed substantial overlap with published E11 AGM HSC signature genes (Vink *et al*., 2020) and adult HSC signature genes, particularly those identified as specific to dormant HSC (Cabezas-Wallscheid *et al*., 2017), compared with Cxcr4-negative HE (Figure S4C, D). Furthermore, among the genes specifically enriched in Cxcr4-positive HE were many with established roles in HSC dormancy and self-renewal, including *H19, Cdkn1c, Ndn* (Necdin), *Mecom*, *Cd81*, *Eng* (Endoglin), and *Igf2* (Asai et al., 2012; Borges et al., 2019; Goyama et al., 2008; Kataoka *et al*., 2011; Lin et al., 2011; Matsumoto *et al*., 2011; Thomas *et al*., 2016; Venkatraman *et al*., 2013). Indeed, expression of these genes, as well as aggregated gene expression of a larger set of published HSC self-renewal/dormancy regulators, demonstrated a similar pattern to arterial EC and HSC signature gene sets, peaking in arterial EC and Cxcr4-positive HE, and significantly decreasing in Cxcr4-negative HE (Figure 4A-B). Notably, expression of HSC self-renewal genes correlates closely with expression of arterial EC genes, as do AGM and adult HSC signature genes, suggesting a close link between early arterial programs and HSC-specifying programs in HSC-competent HE (Figure S4E).

**Figure 4.**
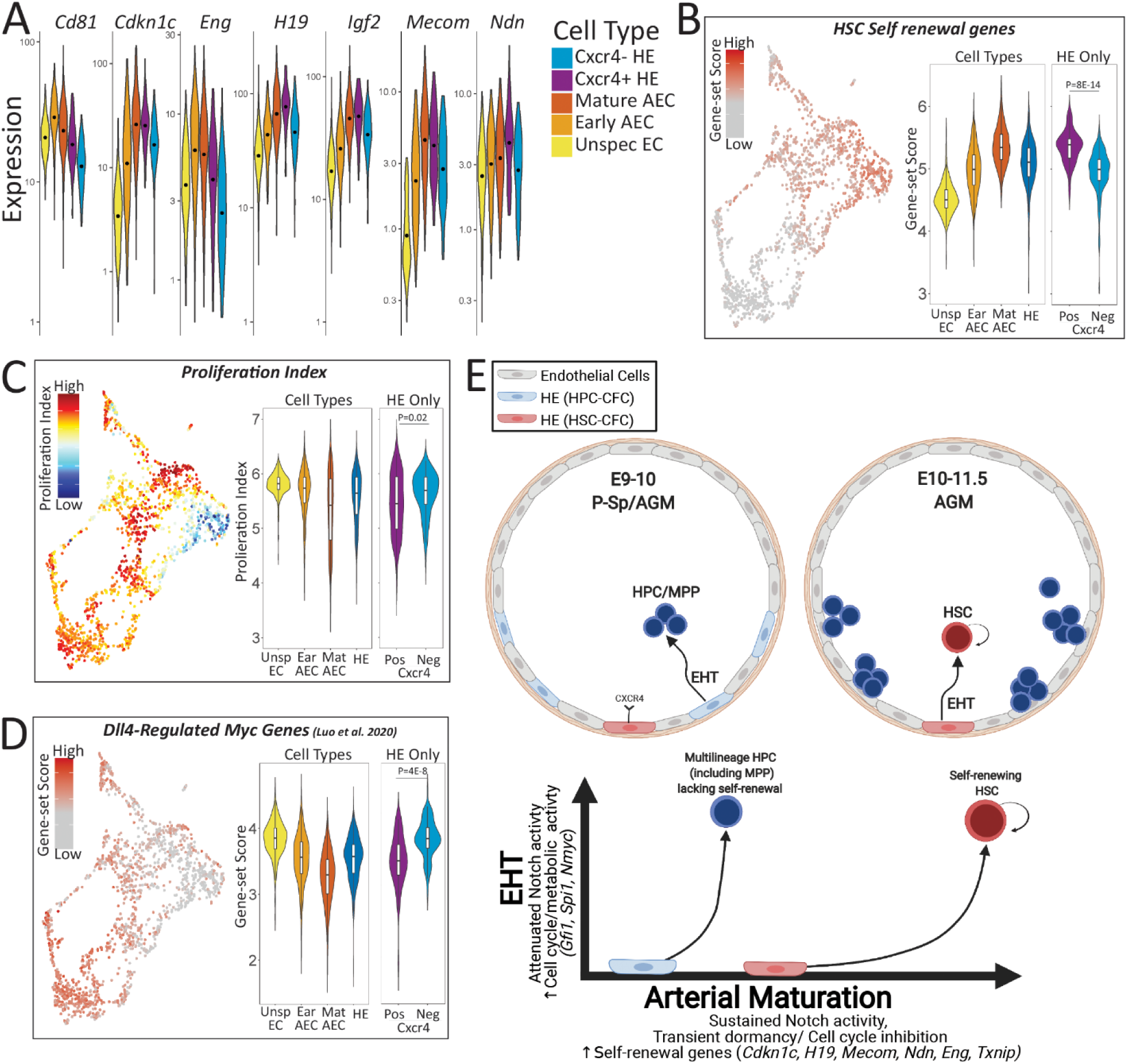
Acquisition of self-renewal programs defining HSC potential. **A.** Violin plots of HSC self-renewal genes identified by differential gene expression between Cxcr4+ and Cxcr4-HE. **B.** UMAP heatmap (left) and violin plots (right) of gene-set scores for published HSC self-renewal genes. **C.** UMAP heatmap (left) and violin plots (right) of proliferation index. **D**. UMAP heatmap (left) and violin plots (right) of gene-set scores for Dll4-regulated Myc pathway genes (Luo *et al*., 2021). **E.** Model depicting distinct waves of multilineage progenitors and HSC derived from HE distinguished by relative expression of genes driving self-renewal tied to arterial maturation and genes driving EHT. Image Created with BioRender.com. (See also Figure S4, Tables S3, S4).

Consistent with the established role of identified HSC self-renewal genes in regulating dormancy by inhibiting cell cycle entry, Cxcr4-positive HE exhibit a significantly lower proliferation index—a transcriptional measure of cell cycle activity— than Cxcr4-negative HE (Figure 4C). Moreover, proliferative status is associated with expression of Myc pathway genes that promote cell cycle and metabolic activity (Luo *et al*., 2021). Cxcr4-negative HE express higher levels of Myc pathway genes that correlate with HE-specific gene expression (Hou *et al*., 2020) (Figure 4D, Figure S4F, G). In contrast, Cxcr4-positive HE express lower levels of Myc genes together with higher Dll4-dependent arterial EC genes and genes associated with dormancy programs (such as embryonic diapause and senescence) (Dhimolea et al., 2021; Duy et al., 2021) (Figure 4D, Figure S4F, G). Altogether, these studies identify the unique molecular signature of HSC-competent HE, supporting a model in which arterial endothelial-associated genes promoting transient dormancy and cell cycle inhibition function as early determinants of HSC fate by establishing self-renewal programs necessary to endow HE with HSC potential (Figure 4E).

## DISCUSSION

In this study, we provide several novel insights into the acquisition of HSC fate during embryonic development. First, we demonstrate that clonal HE with HSC potential can be prospectively distinguished from more abundant HE that lack HSC potential by CXCR4 surface expression, suggesting that cell intrinsic properties necessary for HSC competence are already established at the level of HE in the early P-Sp/AGM. Next, we provide evidence of clonal HE with multilineage hematopoietic progenitor activity that are distinct from HSC-competent HE, identifying an early wave of previously unappreciated HSC-independent MPP activity and suggesting a novel paradigm in which the acquisition of multilineage hematopoietic potential is initially uncoupled from HSC potential in HE at this stage in development (Figure 4E). These findings have important implications for efforts to generate HSC from pluripotent stem cells (PSC), accounting for the observation that PSC-derived HE can clonally generate multilineage hematopoietic progenitors in the absence of HSC potential (Ditadi et al., 2015).

Finally, we used single cell transcriptomics to gain insight into the transcriptional programs that regulate the HSC potential of HE, demonstrating that the Cxcr4-expressing subset of HE is enriched in genes required for HSC self-renewal which are closely associated with arterial transcriptional programs. Notably, we and others have recently shown that the retention of arterial endothelial transcriptional signatures is a key feature that distinguishes pre-HSC and the first engrafting HSC from more differentiated hematopoietic progenitors in the later (E10.5-E11.5) AGM (Hadland *et al*., 2021; Vink *et al*., 2020). Another recent study used single cell approaches to identify a set of genes that distinguish “HSC-primed” HE from non-hemogenic arterial EC in the P-Sp/AGM (Hou *et al*., 2020); however, we found that this set of genes is more highly expressed in Cxcr4-negative HE than in the Cxcr4-positive HE that possess HSC potential. Thus, these genes likely represent early markers and/or drivers of multilineage hematopoiesis during EHT in the E9-9.5 P-Sp/AGM rather than specific markers of HSC fate. In contrast, our data suggest that HSC-competence is uniquely defined by the balanced expression of these HE-specific genes together with HSC self-renewal factors tied to arterial transcriptional programs, implying that the precise timing and dynamic modulation of signals promoting arterial maturation of HE and those driving EHT may be essential to enable HSC fate (Figure 4E).

Our results also suggest that transient dormancy characterized by delayed cell cycle entry and low metabolic activity may be a feature of HSC-competent HE in the early P-Sp/AGM, based on the lower expression of cell cycle and Myc pathway genes, and higher expression of known HSC cell cycle regulators such as *Mecom*, *Necdin*, and *Cdkn1c* (Kataoka *et al*., 2011; Kustikova et al., 2013; Matsumoto *et al*., 2011). This is consistent with studies in Fucci reporter mice, in which HSC precursor activity is restricted to cells in G_0_/G_1_ at E9.5 but is subsequently detected in actively cycling cells in the E10.5-11.5 AGM (Batsivari et al., 2017). Future studies are required to identify a precise mechanism linking metabolic and proliferative dormancy to HSC fate during HE specification. Interestingly, a reversible dormancy state mediated by Myc pathway inhibition has been implicated in embryonic diapause, maintenance of adult HSC, and resistance of cancer cells to chemotherapy, and Myc pathway inhibition was also recently shown to regulate arterial EC fate downstream of Dll4-Notch signaling (Cabezas-Wallscheid *et al*., 2017; Dhimolea *et al*., 2021; Duy *et al*., 2021; Luo *et al*., 2021; Scognamiglio et al., 2016). Notch signaling has also been implicated in arterial EC fate by promoting cell cycle arrest via a GJA4-CDKN1B axis downstream of blood flow (Fang et al., 2017) and CXCL12-CXCR4 signaling is also known to promote adult HSC quiescence (Nie et al., 2008), suggesting additional potential mechanisms. Interestingly, Porcheri and colleagues (Porcheri et al., 2020) recently demonstrated a role for the Notch ligand DLL4 in inhibiting recruitment of HE to nascent intra-aortic hematopoietic clusters in the AGM. Combined with our current studies, this suggests a model in which DLL4-mediated Notch activation could transiently delay EHT from a subset of Cxcr4-expressing HE in the E9-9.5 P-Sp, promoting further maturation of arterial programs that we show are closely associated with expression of HSC self-renewal/dormancy genes. This model would account for the sequential emergence of MPP/HPC activity in the E9-10 P-Sp that lack self-renewal/engraftment properties followed by a burst of pre-HSC/HSC activity peaking between E10.5 and E11.5 (Figure 4E) (Hadland *et al*., 2021; Rybtsov et al., 2016). It further suggests the need for precise temporal modulation of Notch activity and downstream arterial-associated pathways to enable HSC fate, which may be critical for efforts to generate HSC from pluripotent stem cells. Altogether, our studies shift the current paradigm of developmental hematopoiesis by revealing a previously unappreciated wave of MPP that emerges from HE independently of HSC, and further provide new insight into the transcriptional mechanisms of self-renewal acquisition required for HSC fate.

## ACKNOWLEDGEMENTS

Research reported in this publication was supported by the American Society of Hematology Scholar Award and the National Heart, Lung, and Blood Institute (NHLBI) of the National Institutes of Health (NIH) under award number K08HL140143. This research was also supported by the National Institute of Diabetes and Digestive and Kidney Diseases of the NIH under Award Number RC2DK114777. We would like to thank the Fred Hutchinson Flow Cytometry Core for assistance with FACS, and members of the Trapnell, Hadland, Bernstein, and Rafii labs for constructive feedback.

## AUTHOR CONTRIBUTIONS

Conceptualization, B.H., T.D., I.D.B., B.V., S.R., and C.T.; Methodology, B.H., T.D., I.D.B., B.V., S.R., and C.T.; Investigation, T.D., S.D., B.H., B.V., O.W., A.H., D.J., and C.N.; Writing—Original Draft, T.D., B.H.; Writing—Review & Editing, all authors; Funding Acquisition, B.H., I.D.B., S.R., and C.T.; Supervision, B.H.

## STAR METHODS

### KEY RESOURCES TABLE

| REAGENT or RESOURCE | SOURCE | IDENTIFIER |
| --- | --- | --- |
| **Antibodies** |  |  |
| CD201 (EPCR) Anti-mouse Monoclonal Antibody PerCP-eFluor710 Clone eBio1560 | eBioscience | Cat#46-2012-80;<br>RRID:AB_10718383 |
| CD61 Anti-mouse/rat Monoclonal Antibody APC Clone 2C9.G2 | Biolegend | Cat#104316;<br>RRID:AB_2561734 |
| CD144 (VE-cadherin) Anti-mouse Monoclonal Antibody PE-Cyanine7 Clone eBioBV13 | eBioscience | Cat#25-1441-82;<br>RRID:AB_2573402 |
| CD45 Anti-mouse Monoclonal Antibody FITC Clone 30-F11 | eBioscience | Cat#11-0451-85;<br>RRID:AB_465051 |
| CD184 (CXCR4) Anti-mouse Monoclonal Antibody PE Clone 2B11 | Invitrogen | Cat#12-9991-81;<br>RRID:AB_891393 |
| CD41 Anti-mouse Monoclonal Antibody FITC Clone MWReg30 | BD Biosciences | Cat#553848;<br>RRID:AB_395085 |
| CD43 Anti-mouse Monoclonal Antibody FITC Clone eBioR2/60 | eBioscience | Cat#11-0431-82;<br>RRID:AB_465040 |
| CD44 Anti-mouse Monoclonal Antibody APC Clone IM7 | eBioscience | Cat#17-0441-82;<br>RRID:AB_469390 |
| Rat IgG2b kappa Isotype Control Antibody PE Clone A95-1 | BD Pharmingen | Cat#553989;<br>RRID:AB_10049479 |
| Rat IgG1 kappa Isotype Control Antibody FITC Clone R3-34 | BD Pharmingen | Cat#553924;<br>RRID:AB_479706 |
| Rat IgM kappa Isotype Control Antibody FITC Clone eBRM | eBioscience | Cat#11-4341-82;<br>RRID:AB_470017 |
| Rat IgG2b kappa Isotype Control | BD Pharmingen | Cat#553988; |
| Antibody FITC Clone A95-1 |  | RRID:AB_479619 |
| Rat IgG2b kappa Isotype Control Antibody APC Clone A95-1 | BD Pharmingen | Cat#553991;<br>RRID:AB_10050405 |
| DLL4 Anti-mouse Monoclonal Antibody PE Clone HMD4-1 | Biolegend | Cat#130808;<br>RRID:AB_2092983 |
| Armenian Hamster IgG Isotype Control Antibody PE Clone eBio299Arm | eBioscience | Cat#12-4888-81;<br>RRID:AB_470073 |
| Ly-6G and LY6C (Gr-1) Rat Anti-mouse Monoclonal Antibody FITC Clone RB6-8C5 | BD Pharmingen | Cat#553127;<br>RRID:AB_394643 |
| F4/80 Anti-mouse Monoclonal Antibody FITC Clone BM8 | eBioscience | Cat#553127;<br>RRID:AB_2637191 |
| CD45 Anti-mouse Monoclonal Antibody PerCP-Cyanine5.5 Clone 30-F11 | eBioscience | Cat#45-0451-82;<br>RRID:AB_1107002 |
| CD201 (EPCR) Anti-mouse Monoclonal Antibody PE Clone eBio1560 | eBioscience | Cat#12-2012-82;<br>RRID:AB_914317 |
| Ly-6A/E (Sca-1) Anti-mouse Monoclonal Antibody APC Clone D7 | eBioscience | Cat#17-5981-82;<br>RRID:AB_469487 |
| CD93 (AA4.1) Anti-mouse Monoclonal Antibody PE Clone AA4.1 | eBioscience | Cat#12-5892-81;<br>RRID:AB_466017 |
| CD19 Anti-mouse Monoclonal Antibody APC Clone 1D3 | BD Pharmingen | Cat#550992;<br>RRID:AB_398483 |
| CD45R (B220) Ant-mouse Monoclonal Antibody PerCP Clone RAB-6B2 | BD Pharmingen | Cat#553093;<br>RRID:AB_394622 |
| CD11b Anti-mouse Monoclonal Antibody APC-eFluor780 Clone M1/70 | eBioscience | Cat#47-011282;<br>RRID:AB_1603193 |
| CD8a Anti-mouse Monoclonal Antibody PE Clone 53-6.7 | BD Pharmingen | Cat#553033;<br>RRID:AB_394571 |
| CD4 Anti-mouse Monoclonal | BD Pharmingen | Cat#553052; |
| Antibody PerCP Clone RM4-5 |  | RRID:AB_394587 |
| CD25 Anti-mouse Monoclonal<br>Antibody PE-Cyanine7 Clone PC61 | BD Pharmingen | Cat#552880;<br>RRID:AB_394509 |
| Rat IgG2a kappa Isotype Control<br>Antibody PerCP Clone R35-95 | BD Pharmingen | Cat#553933;<br>RRID:AB_10056901 |
| Rat IgG2b kappa Isotype Control<br>Antibody APC-Cy7 Clone A95-1 | BD Pharmingen | Cat#552773;<br>RRID:AB_394459 |
| Rat IgG1 kappa Isotype Control Antibody<br>PE-Cyanine7, Clone eBRG1 | eBioscience | Cat#25-4301-82;<br>RRID:AB_470198 |
| Rat IgG2a kappa Isotype Control<br>Antibody PE Clone eBR2a | eBioscience | Cat#12-4321-80;<br>RRID:AB_1834380 |
| Rat IgG2b kappa Isotype Control<br>Antibody PerCP-Cyanine5.5 Clone RTK4530 | Biolegend | Cat#400632<br>RRID:AB_893677 |
| Rat IgG2a kappa Isotype Control<br>Antibody APC Clone R35-95 | BD Pharmingen | Cat#553932;<br>RRID:AB_479720 |
| Rat IgG2a kappa Isotype Control<br>Antibody APC-eFluor 780 Clone eBR2a | eBioscience | Cat#47-4321-82<br>RRID:AB_11063700 |
| CD45.1 Anti-mouse Monoclonal Antibody<br>PE-Cyanine7 Clone A20 | eBioscience | Cat#25-0453-82<br>RRID:AB_469629 |
| CD45.2 Anti-mouse Monoclonal<br>Antibody APC-eFluor 780 Clone 104 | eBioscience | Cat#47-0454-82<br>RRID:AB_1272175 |
| CD3 Rat Anti-Mouse Monoclonal<br>Antibody FITC Clone 17A2 | BD Pharmingen | Cat#555274<br>RRID:AB_395698 |
| F4/80 Anti-mouse Monoclonal<br>Antibody PE Clone BM8 | Biolegend | Cat#123110<br>RRID:AB_893486 |
| Ly-6G and Ly-6C (Gr-1) Rat Anti-Mouse<br>PerCP-Cyanine5.5 Clone RB6-8C5 | BD Pharmingen | Cat#552093<br>RRID:AB_394334 |
| CD19 Anti-Mouse Monoclonal Antibody<br>APC Clone 1D3 | Biolegend | Cat#152410<br>RRID:AB_2629839 |

| **Bacterial and Virus Strains** |  |  |
| --- | --- | --- |
| Lentivirus containing Myr-Akt |  |  |
| **Chemicals, Peptides, and Recombinant Proteins** |  |  |
| X-Vivo20 | Lonza | Cat#04-448Q |
| Recombinant Murine SCF | Peprotech | Cat#250-03 |
| Recombinant Murine IL-3 | Peprotech | Cat#213-13 |
| Recombinant Human TPO | Preprotech | Cat#300-18 |
| Human Flt3-Ligand | Miltenyi Biotec | Cat#130-096-479 |
| Human IL7 | Miltenyi Biotec | Cat#130-095-363 |
| Fetal Bovine Serum | ThermoFisher | Special order<br>characterized lot |
| Dulbecco's Phosphate Buffered Saline (PBS) | Gibco | Cat#14190-144 |
| Collagenase Type 1 (0.25%) | Stemcell Technologies | Cat#07902 |
| Anti-mouse CD16/32 (FcR block) | BD Biosciences | Cat#553141 |
| MEM alpha | Gibco | Cat# 12561056 |
| Penicillin/StreptoMycin (100X) | Gibco | Cat# 15140-122 |
| TrypLE Express | Gibco | Cat#12605-028 |
| Methocult GF M3434 | Stemcell Technologies | Cat#03434 |
| EmbryoMax 0.1% Gelatin solution | MilliporeSigma | Cat#ES006B |
| DAPI | Millipore | Cat#268298 |
| Iscove's Modified Dulbecco's Medium (IMDM) | Gibco | Cat#12440-053 |
| Heparin | Sigma | Cat# H3149-100KU |
| L-Glutamine | Stemcell Technologies | Cat#7100 |
| Endothelial Mitogen *No longer available—<br>substitute with rm VEGF, rm FGF, rm IGF, | Alfa Aesar | Cat#BT-203 |
| rm EGF |  |  |
| Recombinant Murine VEGF | Peprotech | Cat# 450-32 |
| Recombinant Murine FGF | Peprotech | Cat# 450-33 |
| Recombinant Murine IGF | Peprotech | Cat# 250-19 |
| Recombinant Murine EGF | Peprotech | Cat# 315-09 |
| **Critical Commercial Assays** |  |  |
| Chromium Single Cell 3' GEM, Library & Gel Bead Kit v2 | 10X Genomics | Cat#120237 |
| Chromium Single Cell B Chip Kit | 10X Genomics | Cat#1000073 |
| Chromium i7 Multiplex Kit | 10X Genomics | Cat#120262 |
| Illumina NextSeq 550 high-output kit | Illumina |  |
| 24-well tissue culture plates | Fisher Scientific | Cat#0720084 |
| 96-well tissue culture plates | Fisher Scientific | Cat#0720092 |
| 48-well tissue culture plates | Fisher Scientific | Cat#0877252 |
| **Deposited Data** |  |  |
| Single Cell RNA Seq | This paper |  |
| **Experimental Models: Cell Lines** |  |  |
| AGM-EC | Hadland et al.<br>2015 |  |
| OP9-Dll4 cell line | Zúñiga-Pflücker<br>lab |  |
| OP9-GFP cell line | Zúñiga-Pflücker<br>lab |  |
| **Experimental Models: Organisms/Strains** |  |  |
| Mouse: C57BL/6J | Bred in house<br>(origination:<br>Jackson<br>laboratories) |  |
| Mouse: C57BL/6.SJL-Ly5.1-Pep3b | Bred in house<br>(origination: |  |
|  | Jackson<br>laboratories) |  |
| **Recombinant DNA** |  |  |
| pCCL-PGK myrAkt | Kobayashi et al.<br>2010 |  |
| **Software and Algorithms** |  |  |
| monocle3 (v.3.2.3.0) |  |  |
| ggplot2 (v 3.3.2) |  |  |
| R (version 3.6.1) |  |  |
| FlowJo version 10.7.1 |  |  |
| CellRanger (v 2.1.1) |  |  |
| garnett (v 0.2.15) |  |  |
| ggpubr (v 0.4.0) |  |  |

### RESOURCE AVAILABILITY

#### Lead Contact

Further information and requests for resources and reagents should be directed to and will be fulfilled by the Lead Contact, Brandon Hadland

#### Materials Availability

This study did not generate new unique reagents. We are happy to provide AGM-EC upon request.

#### Data and Code Availability

Raw sequencing data and Monocle 3 cell data sets have been uploaded to NCBI GEO (accession number GSE171457). R scripts used for data analysis are available upon request.

### EXPERIMENTAL MODEL AND SUBJECT DETAILS

#### Mice

Wild type C57Bl6/J7 (CD45.2) and congenic C57BL/6.SJL-Ly5.1-Pep3b (CD45.1) mice were bred at the Fred Hutchinson Cancer Research Center. Male and female C57Bl6/J7 CD45.2 mice at 8-12 weeks of age were used for timed matings and transplantation experiments. All animal studies were conducted in accordance with the NIH guidelines for humane treatment of animals and were approved by the Institutional Animal Care and Use Committee at the Fred Hutchinson Cancer Research Center.

#### Cell Lines

##### AGM-derived Akt-EC (AGM-EC)

AGM-EC were generated as previously described (Hadland *et al*., 2015) and further detailed in a protocol available at Nature Protocol Exchange (https://protocolexchange.researchsquare.com/). For reproducibility, AGM-EC from frozen aliquots of the same derivation were used for all co-culture experiments at less than passage 15 for experiments in this manuscript. AGM-EC were plated at a density of 5×10^5^ cells/flask in gelatinized, tissue culture T75 flasks in EC media (consisting of IMDM with 20% FBS, Penicillin/streptoMycin, Heparin, L-glutamine, and EC mitogen*), and passaged every 3-4 days when confluent.

*EC mitogen no longer available (see STAR Key Resources Table for details on substitutes)

##### OP9/OP9-DLL4 bone marrow stromal cells (OP9 cells)

GFP-expressing OP9 cells and GFP-expressing OP9 stromal cells ectopically expressing Delta-like 4 (OP9-DLL4 cells) generated by the Zúñiga-Pflücker lab (Schmitt and Zúñiga-Pflücker, 2006) were used to assay the B- and T-lymphoid potential of hematopoietic colony cells. OP9 cell lines were plated at a density of 1.6×10^5^ cells/flask in gelatinized, tissue culture T25 flasks in OP9 media (consisting of aMEM with 20% FBS, and Penicillin/streptoMycin). Cells were passaged every 3 days (or when confluent) and used at passage numbers below 13 for co-culture assays.

### METHOD DETAILS

#### Embryo dissections and cell sorting

Embryonic P-Sp/AGM regions were dissected from embryos harvested from pregnant C57Bl6/J7 (CD45.2) female mice as previously described (Hadland *et al*., 2015). Embryo age was precisely timed by counting somite pairs (sp), defined as follows (except where more specifically indicated in the figures): E9 (13-20sp), E9.5 (21-29 sp), and E10 (30-39sp). Dissected P-Sp/AGM tissues were treated with 0.25% collagenase for 25 minutes at 37°C, pipetted to single cell suspension, and washed with PBS containing 10% FBS. Cells were pre-incubated with anti-mouse CD16/CD32 (FcRII block) and stained with monoclonal antibodies as indicated. A detailed list of all antibodies used is shown in the STAR Key Resources Table. For most experiments, to isolate hemogenic endothelial populations, a combination of anti-mouse VE-Cadherin-PECy7, EPCR/CD201-PerCP-eFluor710, and/or CD61-APC were used, with or without additional anti-mouse antibodies CXCR4 (PE), CD41 (FITC), CD43 (FITC), CD45 (FITC), and CD44 (APC), as indicated in the results section. Relevant isotype control antibodies were used to set gates. DAPI staining was used to gate out dead cells. All reagents for cell staining were diluted in PBS with 10% FBS and staining was carried out on ice or at 4°C. Cells were sorted on a BD FACSAria II equipped with BD FACSDiva Software with index sorting capability. For index-sorted single cells, sorting was performed in single cell mode with maximum purity mask settings to minimize contaminating cells.

#### AGM-EC co-culture

For co-culture experiments, AGM-EC at passage 15 or less were plated 24-48 hours prior to initiation of co-culture at a density of 1×10^4^ cells per well to gelatin-treated 96-well tissue culture plates for single cell index co-culture or 4×10^4^ cells per 24-well for bulk co-culture. For single cell index co-culture, AGM-derived hemogenic cells were individually index sorted to each well of 96-well containing AGM-EC in serum-free media consisting of X-vivo 20 with recombinant cytokines: stem cell factor (SCF) and receptor-type tyrosine protein kinase (FLT-3) each at 100 ng/ml, and interleukin-3 (IL-3) and thrombopoietin (TPO) each at 20 ng/ml. Following 6 to 7 days of co-culture, each well was visualized for hematopoietic colony formation and 50% of cells were harvested by vigorous pipetting for phenotypic analysis by flow cytometry, and in some experiments, remaining cells were used for confirmatory transplantation assay as previously described (Hadland *et al*., 2017; Hadland *et al*., 2018) and/or secondary hematopoietic lineage potential assays (CFU assay and OP9 co-culture, described below). For co-culture of bulk populations, sorted cells were re-suspended in serum-free X-vivo 20 culture media with cytokines (excluding FLT-3) and plated at 3-5 embryo equivalent of cells per 24-well containing AGM-EC. Following 6 to 7 days of co-culture, hematopoietic progeny were harvested by vigorous pipetting for subsequent analysis by flow cytometry and transplantation assays, as described below.

#### Flow cytometry analysis of cultured cells

Following co-culture, a fraction of the generated hematopoietic progeny was harvested by vigorous pipetting from the EC layer for analysis of surface phenotype by flow cytometry (approximately 10% per well of cells cultured in bulk on AGM-EC or engineered conditions, or 50% of cells generated following single cell index culture on AGM-EC). Cells were spun and re-suspended in PBS with 2% FBS, pre-incubated with anti-mouse CD16/CD32 (FcRII block) and then stained with the following anti-mouse monoclonal antibodies: VE-Cadherin-PeCy7, CD45-PerCP, Gr-1-FITC, F4/80-FITC, Sca-1-APC, and EPCR-PE. DAPI was used to exclude dead cells. Flow cytometry was performed on a Becton Dickinson Canto 2 and data analyzed using FlowJo Software. Cells with HSC potential were identified phenotypically as VE-cadherin^-/low^ CD45^+^Gr1^-^ F4/80^-^Sca1^high^EPCR^high^. We previously showed that detection of cells with this HSC phenotype following in vitro culture correlated with long-term multilineage engraftment as measured by transplantation assays performed in parallel, whereas hematopoietic progeny without detectable phenotypic HSC did not provide detectable long-term multilineage hematopoietic engraftment (Hadland *et al*., 2017; Hadland *et al*., 2018).

#### Transplantation assays

Following co-culture, a fraction of the generated hematopoietic progeny was harvested by vigorous pipetting from the EC layer, washed with PBS with 2% FBS and re-suspended in 100 ul PBS/2% FBS per mouse transplanted. For bulk co-culture experiments on AGM-EC, the remaining 90% of cells in each well following flow cytometry analysis were pooled, washed with PBS with 2% FBS, and re-suspended at 2-3 embryo equivalent of cells per 100ul PBS/2% FBS, and combined with 5×10^4^ whole marrow cells from adult congenic C57BL/6.SJL-Ly5.1-Pep3b (CD45.1) mice in 100 ul PBS/2% FBS to provide hematopoietic rescue. For some single cell index assays, to validate the presence of functional HSC in colonies containing hematopoietic progeny with phenotypic HSC (VE-cadherin^-/low^CD45^+^Gr1^-^F4/80^-^Sca1^high^EPCR^high^) following co-culture, the remaining 50% volume of each 96-well was harvested for transplantation to individual mice, resuspended in 100ul, and combined with 1×10^5^ CD45.1 whole marrow cells in 100 ul PBS/2% FBS. Cell suspensions (200 ul total volume/mouse) were injected into lethally irradiated (1,000 cGy using a Cesium source) congenic CD45.1 adult recipients via the tail vein. Flow cytometry analysis of peripheral blood obtained by retro-orbital bleeds was performed at various intervals between 2 and 24 weeks following transplantation. Lineage-specific staining for donor (CD45.2) and recipient/rescue (CD45.1) cells from peripheral blood was performed as previously described (Hadland *et al*., 2015), using anti-mouse monoclonal antibodies indicated in the STAR Key Resources Table: CD45.1, CD45.2, CD3, CD19, Gr1, and F4/80. Multilineage engraftment was defined >5% donor (CD45.2) contribution to the peripheral blood with contribution to each lineage of donor myeloid (Gr-1 and F4/80), B cells (CD19) and T-cells (CD3) detected at ≥0.5% at 16 to 24 weeks post-transplant, as indicated.

#### Quantification of HSC and HPC colony-forming cells

Following single cell index co-cultures described above, each individual sorted cell was classified based on its HSC potential. Specifically, single cells giving rise to colonies with detectable HSC by phenotype analysis were categorized as HSC colony-forming cells (HSC-CFC); those giving rise to colonies of CD45^+^ hematopoietic progeny lacking phenotypic HSC were categorized as HPC colony-forming cells (HPC-CFC), and those failing to give rise to detectable hematopoietic colonies were indicated as “no colony.” Results from multiple, pooled index sort experiments from E9.5-E10 (19-30 sp) were combined to quantify the total number of HSC-CFC and HPC-CFC within the VE-Cadherin^+^CD61^+^EPCR^+^ (V^+^E^+^61^+^) sort gate detected per P-Sp/AGM (calculated based on the number of HPC-CFC or HSC-CFC detected, the number of embryo equivalents used, and the fraction of cells in FACS that were index sorted, for each experiment) (Summarized results in Figure 1B).

#### Secondary assays for hematopoietic lineage potential of HPC-CFC

##### CFU assay

Following flow cytometry analysis, a subset of hematopoietic colonies derived from index-sorted HPC-CFC were harvested for colony-forming unit (CFU) assays and OP9 co-culture to evaluate for erythroid and myeloid, and B-and T-lymphoid potential, respectively. In preparation for CFU assays, methylcellulose-based medium with recombinant cytokines was plated to 96-well tissue culture plates. The remaining HPC-CFC progeny following flow cytometry analysis were harvested by vigorous pipetting from the EC layer, washed in serum-free media, and resuspended in serum-free media. One third of the cells from each colony were added to individual 96-wells containing methylcellulose/cytokines (the remaining 2/3 were used for OP9 co-culture described below). Visual characterization of resultant CFU colonies—CFU-Erythroid (CFU-E), CFU-macrophage (CFU-M), CFU-granulocyte (GFU-G), and CFU-granulocyte/macrophage (CFU-GM) was performed under a light microscope for 10-12 days.

##### OP9 co-culture assay

OP9 and OP9 DLL4-expressing (OP9-DLL4) cells were plated 24-48 hours prior to OP9 co-culture initiation at a density of 1×10^4^ cells/well to gelatin-treated 24-well tissue culture plates in OP9 media. On the day of co-culture, 1mL/well of OP9 co-culture media containing FLT-3 and interleukin-7 (IL-7) at 5 ng/ml each was added to OP9 24-wells in place of the OP9 media used for plating. The remaining 2/3 of HPC-CFC progeny processed as described above were divided equally and plated to OP9 or OP9-DLL4 coated wells.

After 6-7 days of co-culture, 10% of cells from the OP9 wells were harvested for flow cytometry analysis. Cells were stained in PBS with 2% FBS containing anti-mouse CD16/32 (FcRII block) and monoclonal antibodies AA4.1 (PE), B220 (Per-CP), CD19 (APC), and CD11b (APC-eFluor780). DAPI was used to exclude dead cells and contaminating GFP-expressing OP9 cells were excluded by gating on the FITC-negative population. Flow cytometry was performed on a Becton Dickinson Canto 2 and data analyzed using FlowJo Software. Isotype control-stained cells were used to set gates for analysis. B-lymphoid cells were identified phenotypically as CD11b^-^ AA4.1^+^CD19^+^B220^+^.

After 8-11 days of co-culture (or when cells began to overgrow), 10% of cells from OP9-DLL4 co-culture were transferred to newly plated OP9-DLL4 wells. After an additional 10-13 days of culture (18-21 days total), 10% of cells from OP9-DLL4 wells were harvested for flow cytometry analysis. Cells were stained in PBS with 2% FBS containing anti-mouse CD16/32 (FcRII block) and monoclonal antibodies CD25 (PE-Cy7), CD44 (APC), CD4 (PerCP), and CD8 (PE). DAPI was used to exclude dead cells, and contaminating GFP-expressing OP9 stromal cells were excluded by gating on the FITC-negative population. Flow cytometry was performed on a Becton Dickinson Canto 2 and data analyzed using FlowJo Software. Isotype control-stained cells were used to set gates for analysis. T-cells were identified phenotypically as CD25^+^CD44^-^CD4^+^CD8^+^.

#### Quantification of progenitor colony-forming cells

Following CFU assays and OP9 co-culture as described above, the individual index-sorted HPC-CFC whose progeny were assessed in secondary assays for hematopoietic lineage potential were further classified according to their particular progenitor potentials. Specifically, HPC-CFC were sub-classified as myeloid progenitor colony-forming cells (Myeloid-CFC), erythromyeloid progenitor colony-forming cells (EMP-CFC), lymphomyeloid progenitor colony-forming cells (LMP-CFC), or multilineage progenitor colony-forming cells (MPP-CFC) according to the combination of myeloid, erythroid, and lymphoid (B-cell and/or T-cell) output that each HPC-CFC gave rise to in CFU and OP9 assays. Cells failing to give rise to detectable hematopoietic colonies and/or hematopoietic output in CFU/OP9 assays were indicated as “no colony/unknown”. Results from multiple, pooled index sort experiments from E9-E10 (19-30 sp) were combined to quantify the total number of Myeloid-CFC, EMP-CFC, LMP-CFC, and MPP-CFC within the VE-Cadherin^+^CD61^+^EPCR^+^ (V^+^E^+^61^+^) sort gate detected per P-Sp/AGM (calculated based on the number of categorized progenitor colonies detected, the number of embryo equivalents used, the fraction of cells in FACS that were index sorted, and the fraction of progenitor colonies that were evaluated in secondary hematopoietic lineage assays, for each experiment) (Summarized results in Figure 2C).

#### Single cell RNA sequencing

For single cell RNA sequencing (scRNA-seq) experiments, freshly sorted AGM-derived cells were washed with PBS containing 0.04% ultrapure BSA and re-suspended in 0.04% ultrapure BSA in PBS on ice. Cell suspensions were loaded into the Chromium Single Cell B Chip (10X Genomics) and processed in the Chromium single cell controller (10X Genomics), targeting a maximum of 3200 cells per lane from freshly sorted AGM-derived cells. The 10X Genomics Version 2 single cell 3’ kit was used to prepare single cell mRNA libraries with the Chromium i7 Multiplex Kit, according to manufacturer protocols. Sequencing was performed for pooled libraries from each sample on an Illumina NextSeq 500 using the 75 cycle, high output kit, targeting a minimum of 50,000 reads per cell. For AGM-derived V^+^61^+^E^+^ cells, scRNA-seq was performed in two independent experiments from cells sorted from embryos pooled at E9 (27 embryos, 12-16 sp) and E9.5 (22 embryos, 15-27 sp), resulting in acquisition of 1843 individual high-quality cell sequences between E9 and E9.5. All sequencing data has been uploaded to NCBI GEO (accession number GSE171457).

#### Single cell transcriptome analysis and quality control

The Cell Ranger 2.1.1 pipeline (10X Genomics) was used to align reads to the mm10 reference genome and generate feature barcode matrix, filtering low-quality cells using default parameters. The Monocle3 (v.3.2.3.0) platform was used for downstream analysis, combining data for cells from each individual sample (as described above) for downstream analysis, using a negative binomial model of distribution with fixed variance, normalizing expression matrices by size factors. Counts for UMI (unique molecular identifiers) and unique genes expressed per cell are shown in boxplots in supplementary figures for each sample (showing median values and interquartile ranges; upper/lower whiskers show 1.5X interquartile range with outliers shown as individual dots). Cells with low UMI counts (<5000), and cells with low genes per cell (<1000) were removed.

#### Dimensionality reduction, batch correction, and cluster analysis

The preprocessCDS() function was used to project the data onto the top principal components (default settings), and the alignCDS() function (Haghverdi et al., 2018) was used to remove batch effects between samples using a “mutual nearest neighbor” algorithm, Batchelor (v.1.2.4). Uniform Manifold Approximation (UMAP) was used for dimensionality reduction with the reduceDimension() function, with reduction_method=’UMAP’. set.seed was used to specify the number of seeds to avoid variability in output due to a random number generator in the function. Clustering was performed with the clusterCells function, with resolution=3E-2 (Figure 3A). Given scattered expression of *Cxcr4* transcript (suggesting low level expression and gene drop-out inherent to scRNAseq data) clusters were classified as Cxcr4-positive or Cxcr4-negative based on threshold detection of *Cxcr4* transcript in >5% of cells in each cluster (Figure 3C), resulting in classification of Cxcr4-positive cells comparable to the expected percentage of CXCR4^+^ cells by flow cytometry (see Figure 1).

#### Cell type classification

Cell type classification was performed using the Garnett package (v.0.2.15) within Monocle 3. Marker gene sets based on established cell type-specific genes (listed below) were used to train a classifier data (using the train_cell_classifier function with default settings) and classify cell types (using the the classify_cells function). Using this method, a small population of cells remains unclassified rather than imputing cell types for all cells in the scRNA-seq dataset. Cell type classifications were used for all downstream analyses.

| **Cell Type** | **Genes Expressed** | **Genes not expressed** |
| --- | --- | --- |
| Unspecified EC | *Etv2*, *Nr2f2* | *Runx1*, *Gfi1*, *Dll4*, *Ptprc* |
| Early arterial EC | *Dll4, Efnb2* | *Runx1, Gfi1, Etv2, Ptprc, Cd44, Gja5* |
| Mature arterial EC | *Cd44, Gja5* | *Runx1, Gfi1, Etv2, Ptprc* |
| HE | *Runx1, Gfi1* | *Etv2, Ptprc* |
| HPC | *Ptprc* |  |

#### Gene set scores

Gene set scores were calculated for each single cell as the log-transformed sum of the size factor-normalized expression for each gene in published signature gene sets or from the Molecular Signatures Database (https://www.gsea-msigdb.org/gsea/msigdb/index.jsp) including: arterial EC marker genes (Aranguren *et al*., 2013; Luo *et al*., 2021; Xu *et al*., 2018), HSC signature genes (Cabezas-Wallscheid *et al*., 2017; Chambers *et al*., 2007; Rodriguez-Fraticelli *et al*., 2020; Wilson *et al*., 2015), HSC self-renewal genes (Asai *et al*., 2012; Ficara et al., 2008; Frelin et al., 2013; Jankovic et al., 2007; Jeong *et al*., 2009; Jude et al., 2007; Kataoka *et al*., 2011; Mallaney et al., 2019; Matsumoto *et al*., 2011; McMahon et al., 2007; Rodriguez-Fraticelli *et al*., 2020; Taoudi et al., 2011; Venkatraman *et al*., 2013; Wang et al., 2016), HE signature genes (Hou *et al*., 2020), AGM HSC genes (Vink 2020), Dll4-regulated Myc target genes (Luo *et al*., 2021), hallmark Myc target genes: https://www.gsea-msigdb.org/gsea/msigdb/cards/HALLMARK_MYC_TARGETS_V1, https://www.gsea-msigdb.org/gsea/msigdb/cards/HALLMARK_MYC_TARGETS_V2, diapause signature genes (Boroviak et al., 2015; Dhimolea *et al*., 2021; Duy *et al*., 2021), senescence signature genes (Duy *et al*., 2021; Fridman and Tainsky, 2008) and cell cycle/proliferation genes (Srivatsan et al., 2020; Tirosh et al., 2016). Violin plots of gene-set scores based on cell types were generated using the ggplot functions geom_violin and geom_boxplot (boxplots show median values and interquartile ranges; upper/lower whiskers show 1.5X interquartile range). For statistical analysis between Cxcr4-negative HE and Cxcr4-postitive HE, Wilcoxon Rank Sum Test (ggupbr package v0.4.0) was used to calculate P values as indicated. Correlation between gene set scores are shown in scatter plots (limited to HE celltype, indicated as Cxcr4-negative HE and Cxcr4-postitive HE) generated using the ggplot functions geom_point and geom_smooth with method set to lm (linear model) to calculate a best fit regression line with 95% confidence intervals indicated by gray shadow.

#### Pseudotime analysis

Pseudotime trajectory analysis was performed using the learnGraph function (with prune_graph=T and ncenter=300 settings to exclude minor trajectories that are not biologically meaningful). The order_cells function was used to calculate where each cell falls in pseudotime. Initial pseudotime was set to cells expressing genes associated with unspecified EC as this represents the most immature cell type in the development hierarchy.

#### Gene module analysis

Gene module analysis was performed using the graph_test functions, which uses a spatial autocorrelation analysis (Moran’s I) that is effective in finding genes that vary in single-cell RNA-seq datasets (Cao *et al*., 2019), and the find_gene_modules function to group genes into modules. For the graph_test function, neighbor_graph was set to “principle graph” to identify modules of genes that co-vary as a function of pseudotime. For the find_gene_modules, resolution was set to 4E-3. Scaled expression scores for gene modules were plotted in UMAP (Figure 3G). Moran’s i and q-values (based on FDR, with FDR<0.05 considered significant) are reported for each gene in the identified gene modules (Table S2). Gene module expression scores were plotted as a function of cluster and cell type to identify modules most specifically associated with regions of UMAP space containing Cxcr4-positive and Cxcr4-negative HE (see Figure S3H).

#### Differential gene expression

Differential gene expression was performed using regression analysis with the fit_models() and coefficient_table() functions in Monocle 3, to identify genes that were differentially expressed based on significance values (q value) adjusted for multiple hypothesis testing using the Benjamini and Hochberg correction method. Adjusted q value <0.05 was considered significant. Cxcr4-positive and Cxcr4-negative HE (by cell type classification) were compared for this analysis, to identify differentially expressed genes in the whole genome (genome-wide, limited to genes expressed in at least 10 cells in the cell data set) (full gene list provided in Table S4). AGM HSC signature genes (Vink *et al*., 2020) and compiled adult HSC signature genes from published data sets (Cabezas-Wallscheid *et al*., 2017; Chambers *et al*., 2007; Rodriguez-Fraticelli *et al*., 2020; Wilson *et al*., 2015) were used identify the subset of genes in Cxcr4-positive vs Cxcr4-negative HE that overlap with HSC signature genes (Figure S4D, C).

#### Gene ontology (GO) analysis

Gene enrichment analysis was performed using two online functional annotations tools, DAVID v6.8 (Huang et al., 2009a; b) or AmiGO v2.5.13 (Ashburner et al., 2000; Carbon et al., 2009; Gene Ontology, 2021). Lists of gene short names for genes found to be significantly upregulated in CXCR4-postive or CXCR4-negative HE were uploaded and mapped to database gene IDs. Functional annotation clustering was performed to identify overrepresented GO-terms for biological processes. Specifically, the ‘GO-FAT’ and ‘GO biological process complete’ algorithms were used for analysis with DAVID and AmiGO, respectively. The top 10 GO-terms from each analysis, as defined by Bonferroni corrected p-values, are reported graphically (Figure S4A, B), while the full analysis can be found in Table S4.

### QUANTIFICATION AND STATISTICAL ANALYSIS

Statistical analyses of colony frequencies were conducted with Prism. Data are expressed as mean +/− standard error (s.e.m), and *n* indicates biological repeats, unless indicated otherwise. P values calculated by student t-test unless otherwise indicated.

## SUPPLEMENTARY INFORMATION

**Supplementary Table 1.** Peripheral blood engraftment of clonal progeny of HSC-CFC and CXCR4+/− bulk sorted populations (Related to Figure 1E, G).

**Supplementary Table 2.** Module-specific genes expressed in modules associated with Cxcr4-positive clusters (module 9) and Cxcr4-negative clusters (module 15) (Related to Figure 3C, G, Figure S3H).

**Supplementary Table 3.** Signature/marker genes from published studies/annotated databases used for gene-set scores. (Related to Fig. 3E, 4B-D, Supplementary Fig. 3G, 4D).

**Supplementary Table 4.** Significantly differentially upregulated genes in Cxcr4-positive HE and Cxcr4-negative HE, and the biological process GO-Terms associated with these gene lists.

---

**Supplementary Figure 1 (Related to Figure 1).**
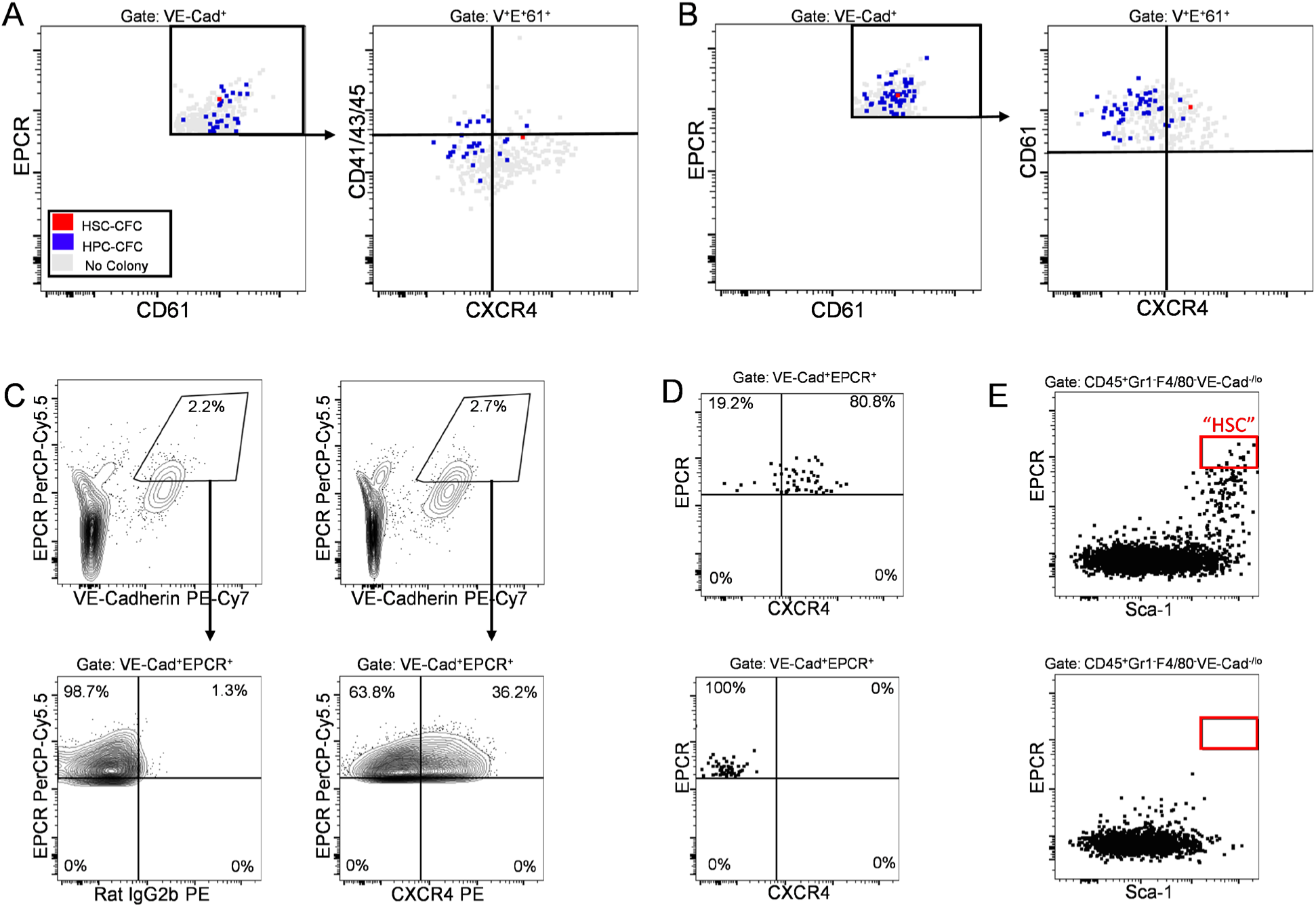
**A-B.** Correlation of clonal HSC-CFC and HPC-CFC with surface expression of CXCR4 and CD41, CD43, and CD45 on individual index-sorted V^+^E^+^61^+^ cells at E9.5. **C.** Gating strategy with isotype controls for CXCR4 (IgG2b PE) within the population sorted as positive for VE-Cadherin and EPCR (left panels), and expression of CXCR4 in the P-Sp/AGM-derived V^+^E^+^ population at E9.5 (22-29 sp)(right panels). **D.** Post-sort purity analysis for CXCR4^+^ (top) and CXCR4^-^ (bottom) bulk-sorted populations from the E9.5 V+E+ population in (A). Gates set according to isotype control. **E.** Flow cytometry for phenotypic HSC following co-culture of sorted CXCR4^+^ (top) and CXCR4^-^ (bottom) populations with AGM-EC.

**Supplementary Figure 2 (Related to Figure 2).**
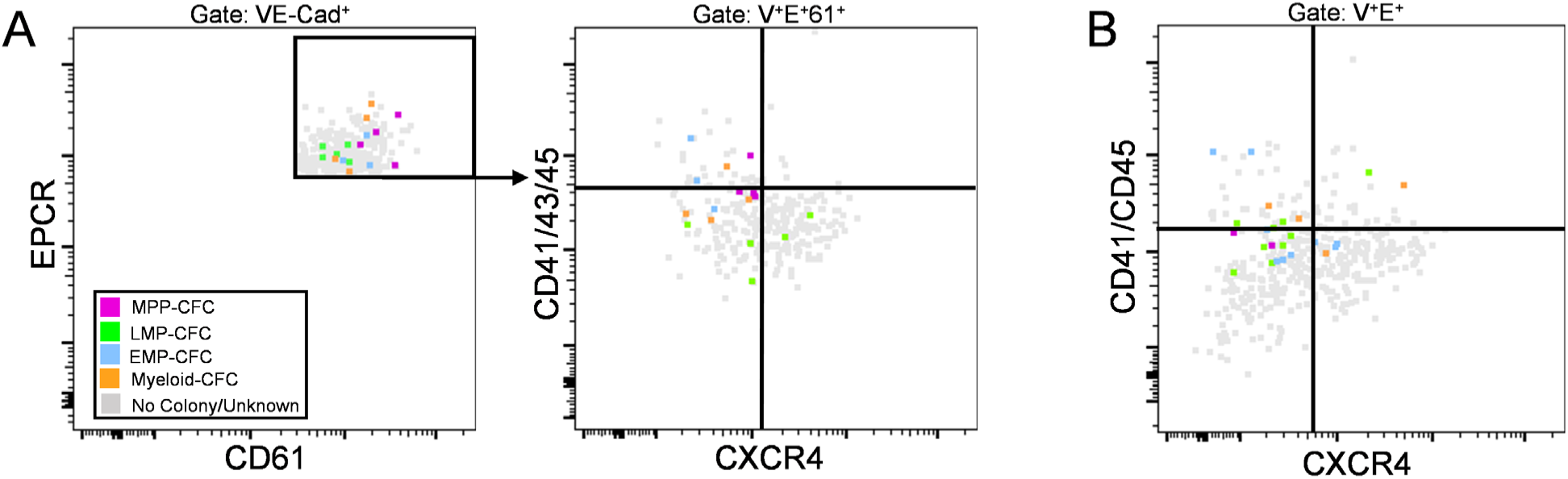
**A-B.** Correlation of clonal hematopoietic lineage potential with CXCR4, CD41, CD43, and CD45 expression of individual HPC-CFC from two independent experiments at E9.5. Multipotent (MPP-CFC), lymphomyeloid (LMP-CFC), erythromyeloid (EMP-CFC), and myeloid colony-forming cells (Myeloid-CFC). No colony/unknown (gray) indicates absence of detectable hematopoietic output following AGM-EC or secondary CFU/OP9 assays.

**Supplementary Figure 3 (Related to figure 3).**
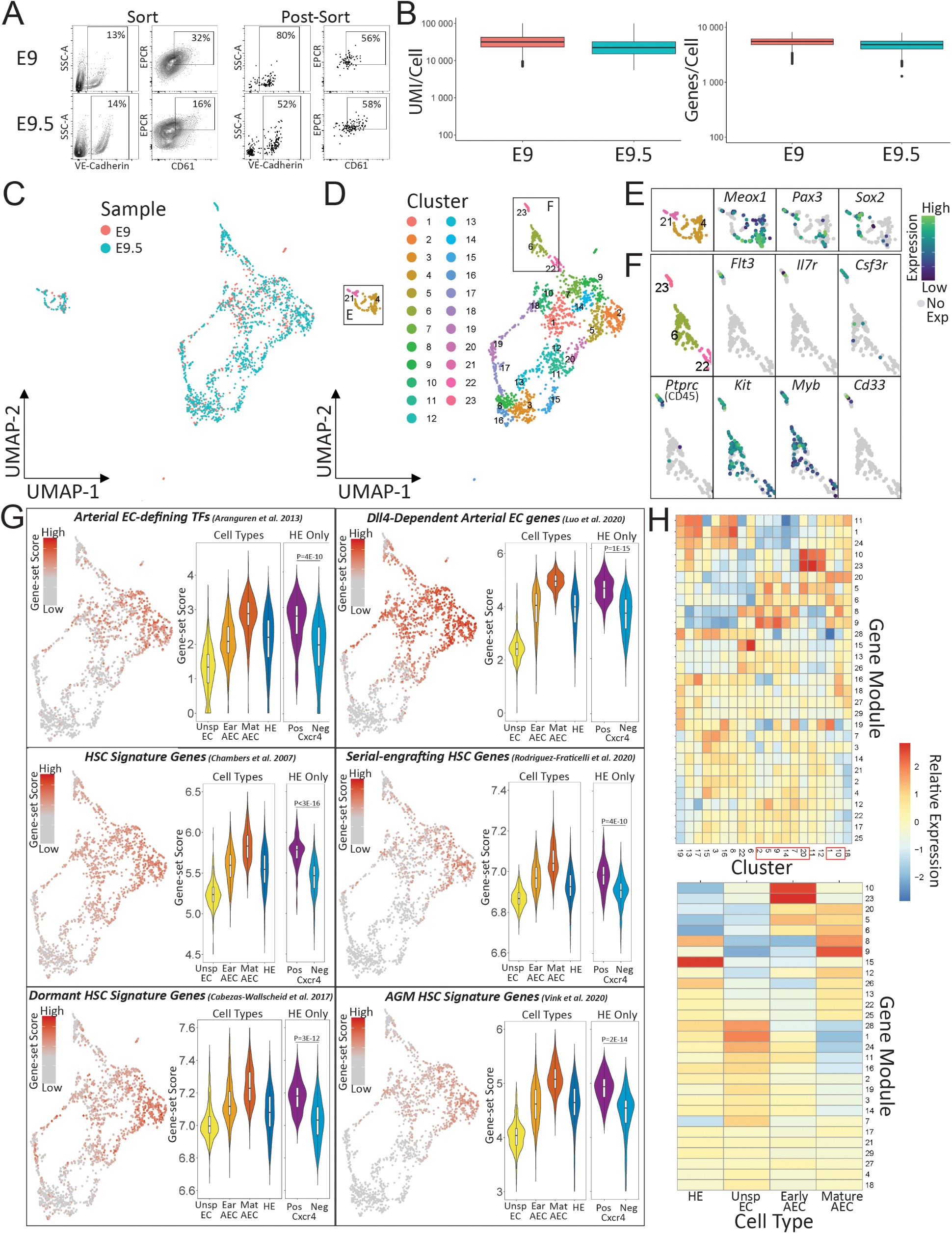
**A.** Sort strategy and post-sort analysis for E9 and E9.5 P-Sp V^+^E^+^61^+^ cells isolated for scRNA-seq. **B.** Counts of UMI and unique genes expressed per cell for E9 and E9.5 P-Sp-derived cell samples after filtering low-quality cells. Boxplots show median values and interquartile ranges; upper/lower whiskers show 1.5X interquartile range with outliers shown as individual dots. **C.** UMAP of 1,841 single cell transcriptomes from 49 embryos, with cells colored by embryo stage/sample. **D.** Unbiased cluster analysis by Leiden community detection. **E-F.** Clusters containing somitic (**E,** clusters 4, 21) and mature HPC (**F,** cluster 23) with expression heatmaps of relevant genes. **G.** Left: Heatmap of gene-set scores for arterial EC-defining transcription factors (Aranguren *et al*., 2013), Dll4-dependent arterial EC genes (Luo *et al*., 2021), HSC signature genes (Chambers *et al*., 2007), serial-engrafting HSC signature genes (Rodriguez-Fraticelli *et al*., 2020), dormant HSC signature gene (Cabezas-Wallscheid *et al*., 2017), and AGM HSC signature genes (Vink *et al*., 2020), Right: Violin plots of gene set scores by cell type, and by Cxcr4+ vs Cxcr4-HE. P values indicate Wilcoxon rank-sum test. **H.** Heatmap of gene expression modules based on cluster (top panel) and cell type (bottom panel). Red outline indicates Cxcr4+ clusters (as defined in Figure 3C).

**Supplementary Figure 4 (Related to Figure 4).**
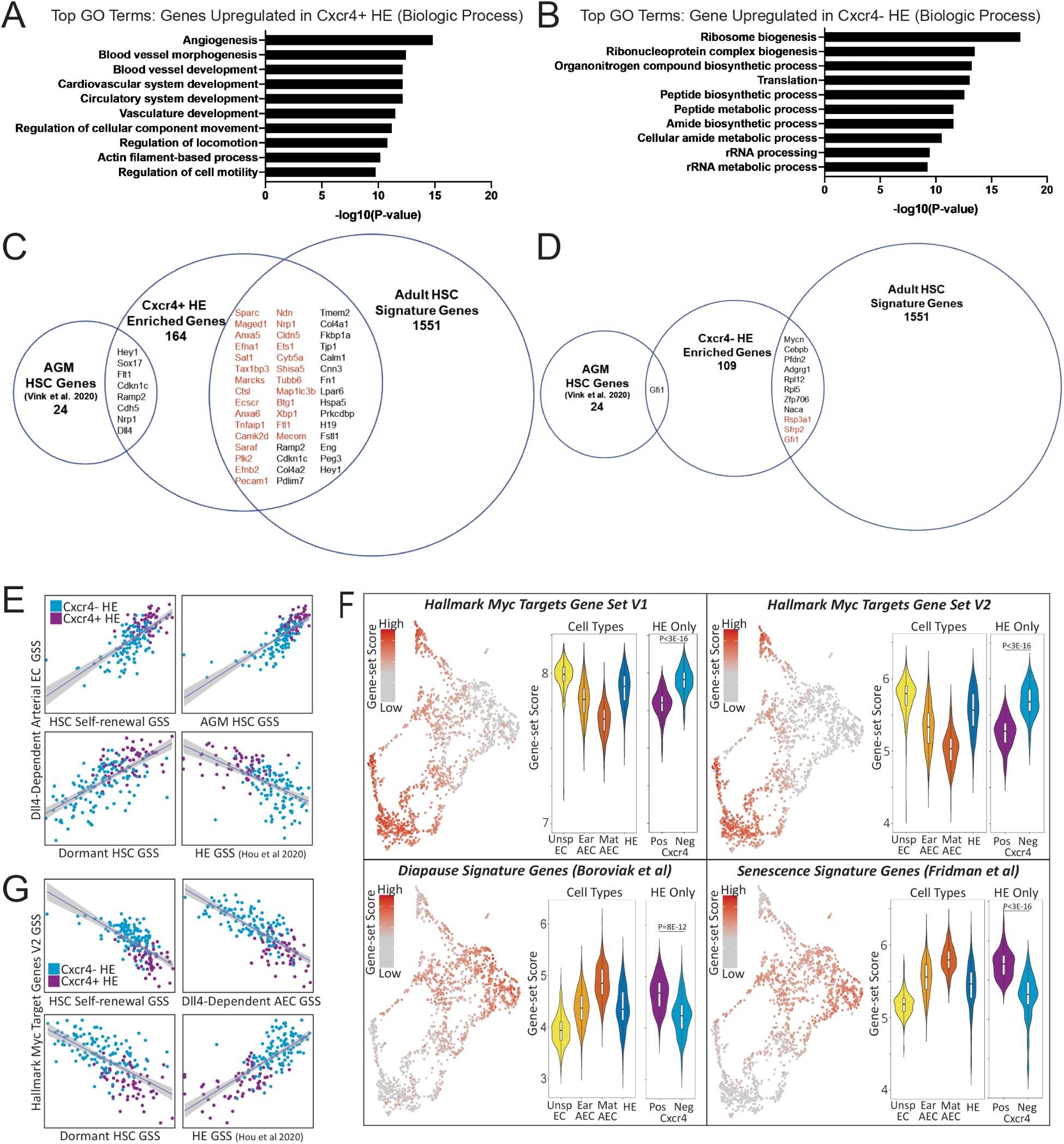
**A-B.** Top gene ontology terms ordered by corrected P value for biological processes amongst genes significantly enriched in Cxcr4-positive (A) vs Cxcr4-negative (B) HE. **C-D.** Genes significantly enriched in Cxcr4-positive HE (C) verses Cxcr4-negative HE (D) that overlap with E11 AGM HSC (Vink *et al*., 2020) or adult HSC signature genes (Cabezas-Wallscheid *et al*., 2017; Chambers *et al*., 2007; Rodriguez-Fraticelli *et al*., 2020; Wilson *et al*., 2015). Genes in red indicate dormant HSC signature gene (Cabezas-Wallscheid *et al*., 2017) **E.** Correlation of gene set-scores for Dll4-dependent arterial EC genes (Luo *et al*., 2021) with gene set scores for HSC self-renewal gene set scores, AGM HSC signature genes (Vink *et al*., 2020), adult dormant HSC signature genes (Cabezas-Wallscheid *et al*., 2017), and HE signature genes (Hou *et al*., 2020) amongst Cxcr4- and Cxcr4+ HE. **F.** Left: Heatmap of gene-set scores for Hallmark Myc Target Genes (Set V1, V2), Diapause signature genes (Boroviak *et al*., 2015), and Senescence signature genes (Fridman and Tainsky, 2008). Right: Violin plots of gene set scores by cell type, and by Cxcr4-positive vs Cxcr4-negative HE. P values indicate Wilcoxon rank-sum test. **G**. Correlation of gene set-scores for Hallmark Myc Target Genes (V2) with gene set scores for HSC self-renewal gene set scores, Dll4-dependent arterial EC genes (Luo *et al*., 2021), adult dormant HSC signature genes (Cabezas-Wallscheid *et al*., 2017), and HE signature genes (Hou *et al*., 2020) amongst Cxcr4- and Cxcr4+ HE.

## REFERENCES

Aranguren, X.L., Agirre, X., Beerens, M., Coppiello, G., Uriz, M., Vandersmissen, I., Benkheil, M., Panadero, J., Aguado, N., Pascual-Montano, A., et al. (2013). Unraveling a novel transcription factor code determining the human arterial-specific endothelial cell signature. Blood 122, 3982–3992. 10.1182/blood-2013-02-483255.

Asai, T., Liu, Y., Di Giandomenico, S., Bae, N., Ndiaye-Lobry, D., Deblasio, A., Menendez, S., Antipin, Y., Reva, B., Wevrick, R., and Nimer, S.D. (2012). Necdin, a p53 target gene, regulates the quiescence and response to genotoxic stress of hematopoietic stem/progenitor cells. Blood 120, 1601–1612. 10.1182/blood-2011-11-393983.

Ashburner, M., Ball, C.A., Blake, J.A., Botstein, D., Butler, H., Cherry, J.M., Davis, A.P., Dolinski, K., Dwight, S.S., Eppig, J.T., et al. (2000). Gene ontology: tool for the unification of biology. The Gene Ontology Consortium. Nat Genet 25, 25–29. 10.1038/75556.

Batsivari, A., Rybtsov, S., Souilhol, C., Binagui-Casas, A., Hills, D., Zhao, S., Travers, P., and Medvinsky, A. (2017). Understanding Hematopoietic Stem Cell Development through Functional Correlation of Their Proliferative Status with the Intra-aortic Cluster Architecture. Stem Cell Reports 8, 1549–1562. 10.1016/j.stemcr.2017.04.003.

Borges, L., Oliveira, V.K.P., Baik, J., Bendall, S.C., and Perlingeiro, R.C.R. (2019). Serial transplantation reveals a critical role for endoglin in hematopoietic stem cell quiescence. Blood 133, 688–696. 10.1182/blood-2018-09-874677.

Boroviak, T., Loos, R., Lombard, P., Okahara, J., Behr, R., Sasaki, E., Nichols, J., Smith, A., and Bertone, P. (2015). Lineage-Specific Profiling Delineates the Emergence and Progression of Naive Pluripotency in Mammalian Embryogenesis. Dev Cell 35, 366–382. 10.1016/j.devcel.2015.10.011.

Böiers, C., Carrelha, J., Lutteropp, M., Luc, S., Green, J.C., Azzoni, E., Woll, P.S., Mead, A.J., Hultquist, A., Swiers, G., et al. (2013). Lymphomyeloid contribution of an immune-restricted progenitor emerging prior to definitive hematopoietic stem cells. Cell Stem Cell 13, 535–548. 10.1016/j.stem.2013.08.012.

Cabezas-Wallscheid, N., Buettner, F., Sommerkamp, P., Klimmeck, D., Ladel, L., Thalheimer, F.B., Pastor-Flores, D., Roma, L.P., Renders, S., Zeisberger, P., et al. (2017). Vitamin A-Retinoic Acid Signaling Regulates Hematopoietic Stem Cell Dormancy. Cell 169, 807–823.e819. 10.1016/j.cell.2017.04.018.

Cao, J., Spielmann, M., Qiu, X., Huang, X., Ibrahim, D.M., Hill, A.J., Zhang, F., Mundlos, S., Christiansen, L., Steemers, F.J., et al. (2019). The single-cell transcriptional landscape of mammalian organogenesis. Nature 566, 496–502. 10.1038/s41586-019-0969-x.

Carbon, S., Ireland, A., Mungall, C.J., Shu, S., Marshall, B., Lewis, S., Ami, G.O.H., and Web Presence Working, G. (2009). AmiGO: online access to ontology and annotation data. Bioinformatics 25, 288–289. 10.1093/bioinformatics/btn615.

Chambers, S.M., Boles, N.C., Lin, K.Y., Tierney, M.P., Bowman, T.V., Bradfute, S.B., Chen, A.J., Merchant, A.A., Sirin, O., Weksberg, D.C., et al. (2007). Hematopoietic fingerprints: an expression database of stem cells and their progeny. Cell Stem Cell 1, 578–591. 10.1016/j.stem.2007.10.003.

de Bruijn, M.F., Speck, N.A., Peeters, M.C., and Dzierzak, E. (2000). Definitive hematopoietic stem cells first develop within the major arterial regions of the mouse embryo. EMBO J 19, 2465–2474. 10.1093/emboj/19.11.2465.

Dhimolea, E., de Matos Simoes, R., Kansara, D., Al’Khafaji, A., Bouyssou, J., Weng, X., Sharma, S., Raja, J., Awate, P., Shirasaki, R., et al. (2021). An Embryonic Diapause-like Adaptation with Suppressed Myc Activity Enables Tumor Treatment Persistence. Cancer Cell 39, 240–256.e211. 10.1016/j.ccell.2020.12.002.

Ditadi, A., Sturgeon, C.M., Tober, J., Awong, G., Kennedy, M., Yzaguirre, A.D., Azzola, L., Ng, E.S., Stanley, E.G., French, D.L., et al. (2015). Human definitive haemogenic endothelium and arterial vascular endothelium represent distinct lineages. Nat Cell Biol 17, 580–591. 10.1038/ncb3161.

Duy, C., Li, M., Teater, M., Meydan, C., Garrett-Bakelman, F.E., Lee, T.C., Chin, C.R., Durmaz, C., Kawabata, K.C., Dhimolea, E., et al. (2021). Chemotherapy induces senescence-like resilient cells capable of initiating AML recurrence. Cancer Discov. 10.1158/2159-8290.cd-20-1375.

Fang, J.S., Coon, B.G., Gillis, N., Chen, Z., Qiu, J., Chittenden, T.W., Burt, J.M., Schwartz, M.A., and Hirschi, K.K. (2017). Shear-induced Notch-Cx37-p27 axis arrests endothelial cell cycle to enable arterial specification. Nat Commun 8, 2149. 10.1038/s41467-017-01742-7.

Ficara, F., Murphy, M.J., Lin, M., and Cleary, M.L. (2008). Pbx1 regulates self-renewal of long-term hematopoietic stem cells by maintaining their quiescence. Cell Stem Cell 2, 484–496. 10.1016/j.stem.2008.03.004.

Frelin, C., Herrington, R., Janmohamed, S., Barbara, M., Tran, G., Paige, C.J., Benveniste, P., Zuñiga-Pflücker, J.C., Souabni, A., Busslinger, M., and Iscove, N.N. (2013). GATA-3 regulates the self-renewal of long-term hematopoietic stem cells. Nat Immunol 14, 1037–1044. 10.1038/ni.2692.

Fridman, A.L., and Tainsky, M.A. (2008). Critical pathways in cellular senescence and immortalization revealed by gene expression profiling. Oncogene 27, 5975–5987. 10.1038/onc.2008.213.

Gene Ontology, C. (2021). The Gene Ontology resource: enriching a GOld mine. Nucleic Acids Res 49, D325–D334. 10.1093/nar/gkaa1113.

Goyama, S., Yamamoto, G., Shimabe, M., Sato, T., Ichikawa, M., Ogawa, S., Chiba, S., and Kurokawa, M. (2008). Evi-1 is a critical regulator for hematopoietic stem cells and transformed leukemic cells. Cell Stem Cell 3, 207–220. 10.1016/j.stem.2008.06.002.

Hadland, B., Varnum-Finney, B., Dozono, S., Dignum, T., Nourigat-McKay, C., Jackson, D.L., Itkin, T., Butler, J.M., Rafii, S., Trapnell, C., and Bernstein, I.D. (2021). Engineering a niche supporting haematopoietic stem cell development using integrated single cell transcriptomics. bioRxiv, 2021.2001.2025.427999. 10.1101/2021.01.25.427999.

Hadland, B., and Yoshimoto, M. (2018). Many layers of embryonic hematopoiesis: new insights into B-cell ontogeny and the origin of hematopoietic stem cells. Exp Hematol 60, 1–9. 10.1016/j.exphem.2017.12.008.

Hadland, B.K., Varnum-Finney, B., Mandal, P.K., Rossi, D.J., Poulos, M.G., Butler, J.M., Rafii, S., Yoder, M.C., Yoshimoto, M., and Bernstein, I.D. (2017). A Common Origin for B-1a and B-2 Lymphocytes in Clonal Pre-Hematopoietic Stem Cells. Stem Cell Reports 8, 1563–1572. 10.1016/j.stemcr.2017.04.007.

Hadland, B.K., Varnum-Finney, B., Nourigat-Mckay, C., Flowers, D., and Bernstein, I.D. (2018). Clonal Analysis of Embryonic Hematopoietic Stem Cell Precursors Using Single Cell Index Sorting Combined with Endothelial Cell Niche Co-culture. J Vis Exp. 10.3791/56973.

Hadland, B.K., Varnum-Finney, B., Poulos, M.G., Moon, R.T., Butler, J.M., Rafii, S., and Bernstein, I.D. (2015). Endothelium and NOTCH specify and amplify aorta-gonad-mesonephros-derived hematopoietic stem cells. J Clin Invest 125, 2032–2045. 10.1172/JCI80137.

Haghverdi, L., Lun, A.T.L., Morgan, M.D., and Marioni, J.C. (2018). Batch effects in single-cell RNA-sequencing data are corrected by matching mutual nearest neighbors. Nat Biotechnol 36, 421–427. 10.1038/nbt.4091.

Hou, S., Li, Z., Zheng, X., Gao, Y., Dong, J., Ni, Y., Wang, X., Li, Y., Ding, X., Chang, Z., et al. (2020). Embryonic endothelial evolution towards first hematopoietic stem cells revealed by single-cell transcriptomic and functional analyses. Cell Res 30, 376–392. 10.1038/s41422-020-0300-2.

Huang, d.W., Sherman, B.T., and Lempicki, R.A. (2009a). Bioinformatics enrichment tools: paths toward the comprehensive functional analysis of large gene lists. Nucleic Acids Res 37, 1–13. 10.1093/nar/gkn923.

Huang, d.W., Sherman, B.T., and Lempicki, R.A. (2009b). Systematic and integrative analysis of large gene lists using DAVID bioinformatics resources. Nat Protoc 4, 44–57. 10.1038/nprot.2008.211.

Inlay, M.A., Serwold, T., Mosley, A., Fathman, J.W., Dimov, I.K., Seita, J., and Weissman, I.L. (2014). Identification of multipotent progenitors that emerge prior to hematopoietic stem cells in embryonic development. Stem Cell Reports 2, 457–472. 10.1016/j.stemcr.2014.02.001.

Jankovic, V., Ciarrocchi, A., Boccuni, P., DeBlasio, T., Benezra, R., and Nimer, S.D. (2007). Id1 restrains myeloid commitment, maintaining the self-renewal capacity of hematopoietic stem cells. Proc Natl Acad Sci U S A 104, 1260–1265. 10.1073/pnas.0607894104.

Jeong, M., Piao, Z.H., Kim, M.S., Lee, S.H., Yun, S., Sun, H.N., Yoon, S.R., Chung, J.W., Kim, T.D., Jeon, J.H., et al. (2009). Thioredoxin-interacting protein regulates hematopoietic stem cell quiescence and mobilization under stress conditions. J Immunol 183, 2495–2505. 10.4049/jimmunol.0804221.

Jude, C.D., Climer, L., Xu, D., Artinger, E., Fisher, J.K., and Ernst, P. (2007). Unique and independent roles for MLL in adult hematopoietic stem cells and progenitors. Cell Stem Cell 1, 324–337. 10.1016/j.stem.2007.05.019.

Kataoka, K., Sato, T., Yoshimi, A., Goyama, S., Tsuruta, T., Kobayashi, H., Shimabe, M., Arai, S., Nakagawa, M., Imai, Y., et al. (2011). Evi1 is essential for hematopoietic stem cell self-renewal, and its expression marks hematopoietic cells with long-term multilineage repopulating activity. J Exp Med 208, 2403–2416. 10.1084/jem.20110447.

Kustikova, O.S., Schwarzer, A., Stahlhut, M., Brugman, M.H., Neumann, T., Yang, M., Li, Z., Schambach, A., Heinz, N., Gerdes, S., et al. (2013). Activation of Evi1 inhibits cell cycle progression and differentiation of hematopoietic progenitor cells. Leukemia 27, 1127–1138. 10.1038/leu.2012.355.

Lin, K.K., Rossi, L., Boles, N.C., Hall, B.E., George, T.C., and Goodell, M.A. (2011). CD81 is essential for the re-entry of hematopoietic stem cells to quiescence following stress-induced proliferation via deactivation of the Akt pathway. PLoS Biol 9, e1001148. 10.1371/journal.pbio.1001148.

Luo, W., Garcia-Gonzalez, I., Fernández-Chacón, M., Casquero-Garcia, V., Sanchez-Muñoz, M.S., Mühleder, S., Garcia-Ortega, L., Andrade, J., Potente, M., and Benedito, R. (2021). Arterialization requires the timely suppression of cell growth. Nature 589, 437–441. 10.1038/s41586-020-3018-x.

Mallaney, C., Ostrander, E.L., Celik, H., Kramer, A.C., Martens, A., Kothari, A., Koh, W.K., Haussler, E., Iwamori, N., Gontarz, P., et al. (2019). Kdm6b regulates context-dependent hematopoietic stem cell self-renewal and leukemogenesis. Leukemia 33, 2506–2521. 10.1038/s41375-019-0462-4.

Matsumoto, A., Takeishi, S., Kanie, T., Susaki, E., Onoyama, I., Tateishi, Y., Nakayama, K., and Nakayama, K.I. (2011). p57 is required for quiescence and maintenance of adult hematopoietic stem cells. Cell Stem Cell 9, 262–271. 10.1016/j.stem.2011.06.014.

McMahon, K.A., Hiew, S.Y., Hadjur, S., Veiga-Fernandes, H., Menzel, U., Price, A.J., Kioussis, D., Williams, O., and Brady, H.J. (2007). Mll has a critical role in fetal and adult hematopoietic stem cell self-renewal. Cell Stem Cell 1, 338–345. 10.1016/j.stem.2007.07.002.

Medvinsky, A., and Dzierzak, E. (1996). Definitive hematopoiesis is autonomously initiated by the AGM region. Cell 86, 897–906.

Müller, A.M., Medvinsky, A., Strouboulis, J., Grosveld, F., and Dzierzak, E. (1994). Development of hematopoietic stem cell activity in the mouse embryo. Immunity 1, 291–301.

Nie, Y., Han, Y.C., and Zou, Y.R. (2008). CXCR4 is required for the quiescence of primitive hematopoietic cells. J Exp Med 205, 777–783. 10.1084/jem.20072513.

Oatley, M., Bölükbası, Ö., Svensson, V., Shvartsman, M., Ganter, K., Zirngibl, K., Pavlovich, P.V., Milchevskaya, V., Foteva, V., Natarajan, K.N., et al. (2020). Single-cell transcriptomics identifies CD44 as a marker and regulator of endothelial to haematopoietic transition. Nat Commun 11, 586. 10.1038/s41467-019-14171-5.

Palis, J. (2016). Hematopoietic stem cell-independent hematopoiesis: emergence of erythroid, megakaryocyte, and myeloid potential in the mammalian embryo. FEBS Lett 590, 3965–3974. 10.1002/1873-3468.12459.

Pliner, H.A., Shendure, J., and Trapnell, C. (2019). Supervised classification enables rapid annotation of cell atlases. Nat Methods 16, 983–986. 10.1038/s41592-019-0535-3.

Porcheri, C., Golan, O., Calero-Nieto, F.J., Thambyrajah, R., Ruiz-Herguido, C., Wang, X., Catto, F., Guillén, Y., Sinha, R., González, J., et al. (2020). Notch ligand Dll4 impairs cell recruitment to aortic clusters and limits blood stem cell generation. EMBO J 39, e104270. 10.15252/embj.2019104270.

Rodriguez-Fraticelli, A.E., Weinreb, C., Wang, S.W., Migueles, R.P., Jankovic, M., Usart, M., Klein, A.M., Lowell, S., and Camargo, F.D. (2020). Single-cell lineage tracing unveils a role for TCF15 in haematopoiesis. Nature 583, 585–589. 10.1038/s41586-020-2503-6.

Schmitt, T.M., and Zúñiga-Pflücker, J.C. (2006). T-cell development, doing it in a dish. Immunol Rev 209, 95–102. 10.1111/j.0105-2896.2006.00353.x.

Scognamiglio, R., Cabezas-Wallscheid, N., Thier, M.C., Altamura, S., Reyes, A., Prendergast Á, M., Baumgärtner, D., Carnevalli, L.S., Atzberger, A., Haas, S., et al. (2016). Myc Depletion Induces a Pluripotent Dormant State Mimicking Diapause. Cell 164, 668–680. 10.1016/j.cell.2015.12.033.

Srivatsan, S.R., McFaline-Figueroa, J.L., Ramani, V., Saunders, L., Cao, J., Packer, J., Pliner, H.A., Jackson, D.L., Daza, R.M., Christiansen, L., et al. (2020). Massively multiplex chemical transcriptomics at single-cell resolution. Science 367, 45–51. 10.1126/science.aax6234.

Taoudi, S., Bee, T., Hilton, A., Knezevic, K., Scott, J., Willson, T.A., Collin, C., Thomas, T., Voss, A.K., Kile, B.T., et al. (2011). ERG dependence distinguishes developmental control of hematopoietic stem cell maintenance from hematopoietic specification. Genes Dev 25, 251–262. 10.1101/gad.2009211.

Thomas, D.D., Sommer, A.G., Balazs, A.B., Beerman, I., Murphy, G.J., Rossi, D., and Mostoslavsky, G. (2016). Insulin-like growth factor 2 modulates murine hematopoietic stem cell maintenance through upregulation of p57. Exp Hematol 44, 422–433.e421. 10.1016/j.exphem.2016.01.010.

Tirosh, I., Izar, B., Prakadan, S.M., Wadsworth, M.H., 2nd, Treacy, D., Trombetta, J.J., Rotem, A., Rodman, C., Lian, C., Murphy, G., et al. (2016). Dissecting the multicellular ecosystem of metastatic melanoma by single-cell RNA-seq. Science 352, 189–196. 10.1126/science.aad0501.

Trapnell, C., Cacchiarelli, D., Grimsby, J., Pokharel, P., Li, S., Morse, M., Lennon, N.J., Livak, K.J., Mikkelsen, T.S., and Rinn, J.L. (2014). The dynamics and regulators of cell fate decisions are revealed by pseudotemporal ordering of single cells. Nat Biotechnol 32, 381–386. 10.1038/nbt.2859.

Venkatraman, A., He, X.C., Thorvaldsen, J.L., Sugimura, R., Perry, J.M., Tao, F., Zhao, M., Christenson, M.K., Sanchez, R., Yu, J.Y., et al. (2013). Maternal imprinting at the H19-Igf2 locus maintains adult haematopoietic stem cell quiescence. Nature 500, 345–349. 10.1038/nature12303.

Vink, C.S., Calero-Nieto, F.J., Wang, X., Maglitto, A., Mariani, S.A., Jawaid, W., Göttgens, B., and Dzierzak, E. (2020). Iterative Single-Cell Analyses Define the Transcriptome of the First Functional Hematopoietic Stem Cells. Cell Rep 31, 107627. 10.1016/j.celrep.2020.107627.

Wang, H., Diao, D., Shi, Z., Zhu, X., Gao, Y., Gao, S., Liu, X., Wu, Y., Rudolph, K.L., Liu, G., et al. (2016). SIRT6 Controls Hematopoietic Stem Cell Homeostasis through Epigenetic Regulation of Wnt Signaling. Cell Stem Cell 18, 495–507. 10.1016/j.stem.2016.03.005.

Wilson, N.K., Kent, D.G., Buettner, F., Shehata, M., Macaulay, I.C., Calero-Nieto, F.J., Sánchez Castillo, M., Oedekoven, C.A., Diamanti, E., Schulte, R., et al. (2015). Combined Single-Cell Functional and Gene Expression Analysis Resolves Heterogeneity within Stem Cell Populations. Cell Stem Cell 16, 712–724. 10.1016/j.stem.2015.04.004.

Xu, C., Gao, X., Wei, Q., Nakahara, F., Zimmerman, S.E., Mar, J., and Frenette, P.S. (2018). Stem cell factor is selectively secreted by arterial endothelial cells in bone marrow. Nat Commun 9, 2449. 10.1038/s41467-018-04726-3.

Zhu, Q., Gao, P., Tober, J., Bennett, L., Chen, C., Uzun, Y., Li, Y., Howell, E.D., Mumau, M., Yu, W., et al. (2020). Developmental trajectory of prehematopoietic stem cell formation from endothelium. Blood 136, 845–856. 10.1182/blood.2020004801.

